# Co-Expression Networks in the Green Alga *Chlamydomonas reinhardtii* Empower Gene Discovery and Functional Exploration

**DOI:** 10.1101/2020.10.05.326611

**Authors:** Patrice A. Salomé, Sabeeha S. Merchant

## Abstract

The unicellular green alga *Chlamydomonas reinhardtii* is a choice reference system for the study of photosynthesis, cilium assembly and function, lipid and starch metabolism and metal homeostasis. Despite decades of research, the functions of thousands of genes remain largely unknown, and new approaches are needed to categorically assign genes to cellular pathways. Growing collections of transcriptome and proteome data now allow a systematic approach based on integrative co-expression analysis. We used a dataset comprising 518 deep transcriptome samples derived from 58 independent experiments to identify potential co-expression relationships between genes. We visualized co-expression potential with the R package *corrplot*, to easily assess co-expression and anti-correlation between genes from manually-curated and community-generated gene lists. We extracted 400 high-confidence cilia-related genes at the intersection of multiple co-expressed lists, illustrating the power of our simple method. Surprisingly, Chlamydomonas experiments did not cluster according to an obvious pattern, suggesting an underappreciated variable during sample collection. One possible source of variation may stem from the strong clustering of nuclear genes as a function of their diurnal phase, even in samples collected in constant conditions, indicating substantial residual synchronization in batch cultures. We provide a step-by-step guide into the analysis of co-expression across Chlamydomonas transcriptome datasets to help foster gene function discovery.

**One-sentence summary:** we reveal co-expression potential between Chlamydomonas genes and describe strong synchronization of cells grown in batch cultures as a possible source of underappreciated variation.

## INTRODUCTION

Discovering the functions of genes has driven biology for over a century, using a multitude of tools to determine the factors associated with a given cellular process. The concept of the gene as a heritable structure was developed by observing how individuals with distinct visible phenotypes could arise from a population and be transmitted to their progeny. Thomas Morgan isolated the first spontaneous mutant in the fruit fly (*Drosophila melanogaster*) in 1910 (Morgan, 1910), followed quickly by more spontaneous mutations through the careful examination of thousands of flies (Bridges and Morgan, 1916). Induced mutations, first by X- or gamma rays, paved the way to classical genetic screens in multiple species, including the Jimson weed (*Datura stramonium*), the fruit fly, the green unicellular alga Chlamydomonas (*Chlamydomonas reinhardtii*) and barley (*Hordeum vulgare*), the latter creating the field of radiation breeding (Gager and Blakeslee, 1927; Muller, 1928; Stadler, 1928; Birch et al., 1953).

These mutations fueled a very thorough phenotypic dissection of the processes affected by the absence of a gene product, but it is only in the 1970s that the nature of the mutated genes began to be unraveled. The development of transformation protocols to introduce transgenes into model systems further opened new possibilities for dissecting the role of a gene in situ by over-expression of a wild-type or mutated copy (Leutwiler et al., 1986; Hinnen et al., 1978; Rubin and Spradling, 1982; Kindle et al., 1989; Rochaix et al., 1984).

The next technological innovation revolutionized biology: deep sequencing techniques have revealed the complete genomic landscape and gene complement of most any species. Expression profiling by microarrays, and later by deep sequencing of the transcriptome (RNAseq) now provide easy access to the changes in the transcriptome in response to genetic or environmental perturbations. In Chlamydomonas alone, RNAseq analysis has empowered hypothesis generation by providing a detailed picture of the changes in gene expression in response to light (Zhu et al., 2008; Xiang et al., 2001; Wittkopp et al., 2017), CO_2_ (Fang et al., 2012; Fukuzawa et al., 2001; Xiang et al., 2001; Brueggeman et al., 2012) and stress (Wakao et al., 2014; Urzica et al., 2012a; Blaby-Haas et al., 2016; Blaby et al., 2015), as well as nutritional deficiencies such as for nitrogen or micronutrients, including iron (Blaby et al., 2013; Castruita et al., 2011; Dudley Page et al., 2012; González-Ballester et al., 2010; Kajikawa et al., 2015; Miller et al., 2010; Ngan et al., 2015; Schmollinger et al., 2014; Urzica et al., 2012b). RNAseq data have largely been analyzed in comparative mode, that is by comparing the wild type to the mutant, or between untreated and treated cultures. Chlamydomonas transcriptome studies comprise hundreds of samples from dozens of independent experiments from multiple research groups. Due to the ease of growing large volumes of cell cultures under defined conditions, While most samples are typically collected from cells grown in constant light, several studies have focused on the diurnal control of gene expression by measuring transcript levels over the course of a diurnal cycle (Strenkert et al., 2019a; Zones et al., 2015; Panchy et al., 2014).

Several pipelines have been implemented that combine transcriptomics datasets to build gene regulatory networks and assign gene function (Romero-Campero et al., 2016; Aoki et al., 2016; Nguyen et al., 2019), based on the premise that genes involved in a similar process will be co-expressed, in particular if their encoded proteins physically interact (Zhu et al., 2008; Simonis et al., 2004; Komurov and White, 2007; Ge et al., 2001). However, these approaches largely allow the visualization of the network associated with a single gene at a time or offer pre-computed co-expression modules; thus, they do not provide a visual summary of the underlying correlations. In addition, negative correlations are not considered. Rather than superseding the contribution of these previous studies, we wished to develop an easily searchable dataset of co-expression and anti-correlation estimates for any gene of interest to facilitate prioritization of candidate genes fulfilling user-defined criteria.

We describe here a thorough analysis of the Chlamydomonas transcriptome landscape, based on the analysis of Pearson’s correlation coefficients associated with all nuclear gene pairs using a set of 518 RNAseq samples from 58 independent experiments. RNAseq samples from a given experiment were more correlated than to samples from any other experiment, even those querying the same variable, indicating the strong environmental sensitivity of Chlamydomonas cultures. We observed frequent co-expression between genes, but also report on anti-correlations, an underappreciated dimension in regulatory networks. We illustrate our approach by revisiting gene lists curated by the Chlamydomonas community and by exploring co-expression modules with the R package *corrplot* (Wei and Simko, 2017) and identify high-confidence candidate genes involved in cilia biogenesis and function. Finally, we discovered that the vast majority of RNAseq samples exhibit substantial diurnal rhythmicity, even when derived from cells grown in constant light. We provide a simple R script for data exploration and hope that this resource will be of use to the community, as this approach can be applied to any biological system.

## RESULTS

### Remapping and Normalization Steps of the Chlamydomonas Transcriptome

The analysis of changes in gene expression typically covers a limited number of conditions on selected genotypes to identify treatment-specific modulators of the transcriptome in a given organism. While this approach is powerful, we wished to integrate multiple transcriptome datasets that represent multiple variables in growth conditions and genotypes. To this end, we collected 58 transcriptome deep-sequencing (RNAseq) datasets generated by the community and by our own laboratory that correspond to 518 samples. We remapped all reads to version v5.5 of the Chlamydomonas genome to remove changes in gene models between experiments as a variable, as our collection of datasets span about 10 years. We did not attempt to compensate for batch effects or variation in sequencing platforms, which were all Illumina-based.

We then assessed the global expression of all 17,741 Chlamydomonas nuclear genes across our set of 518 samples. Most nuclear genes were expressed at levels of 1 Fragment Per Kilobase of transcript per Million mapped reads (FPKM) in the majority of samples, although a large subset of nuclear genes was seldom expressed even at this low expression cut-off (Supplemental Table 1). With a higher threshold for expression, the fraction of expressed nuclear genes decreased (Supplemental Table 1). This pattern indicated that most genes are expressed at moderate levels and only in a limited number of conditions.

We next normalized our RNAseq dataset following the same steps used for the ALCOdb gene co-expression database for microalgae (illustrated in Supplemental Figure 1; Aoki et al., 2016). The final normalization step centered expression estimates to zero, as a Z-score normalization would (Supplemental Figure 1B). *RIBOSOMAL PROTEIN GENES* (*RPG*s) beautifully illustrated the power of normalization (Supplemental Figure 2). Indeed, variation between *RPG*s only emerged after log_2_ normalization, but offered little differentiation on the basis of experiments or samples. Normalization to mean fixed this issue, and revealed variation between *RPGs* and experimental samples that were until then hidden.

### Samples from the Same Experiment Show Strong Positive Correlations

These datasets allowed us to assess the extent of correlation between samples/experiments (each sample being represented by its unique 17,741 gene expression estimates) or between genes (each gene being characterized by its unique 518 gene expression estimates across all samples). We used the R package *corrplot* to visualize correlations across samples or genes (see Supplemental Figure 3 for details). FPKM values failed to extract a pattern, as most samples were strongly and positively correlated, based on Pearson’s correlation coefficients (PCCs) between samples (Figure 1A; mean PCC = 0.74 ± 0.18). The same held true for log_2_- and quantile-normalized datasets (Supplemental Figure 4; mean PCC of 0.83 ± 0.17). It was only after normalization to means that clear localized correlation clusters appeared along the diagonal of the matrix that matched with each experiment (Figure 1B). Indeed, although the entire correlation matrix had a mean PCC close to zero (0.002 ± 0.226), samples belonging to the same experiment exhibited strong and positive correlations (Figure 1C). Samples from a given experiment (including the reference or control samples) were more related to each other than to any other sample, even when designed to query the same biological question (see, for example, nitrogen deprivation samples, Figure 1C and Supplemental Figure 4E). Likewise, the laboratory provenance of samples did not explain the extent of relationship between samples: over half of all RNA-seq samples analyzed here have been generated by our laboratory, and yet most failed to exhibit significant correlations outside of each experiment (Supplemental Figure 4F), despite careful considerations of consistent sample collection procedures.

**Figure 1.**
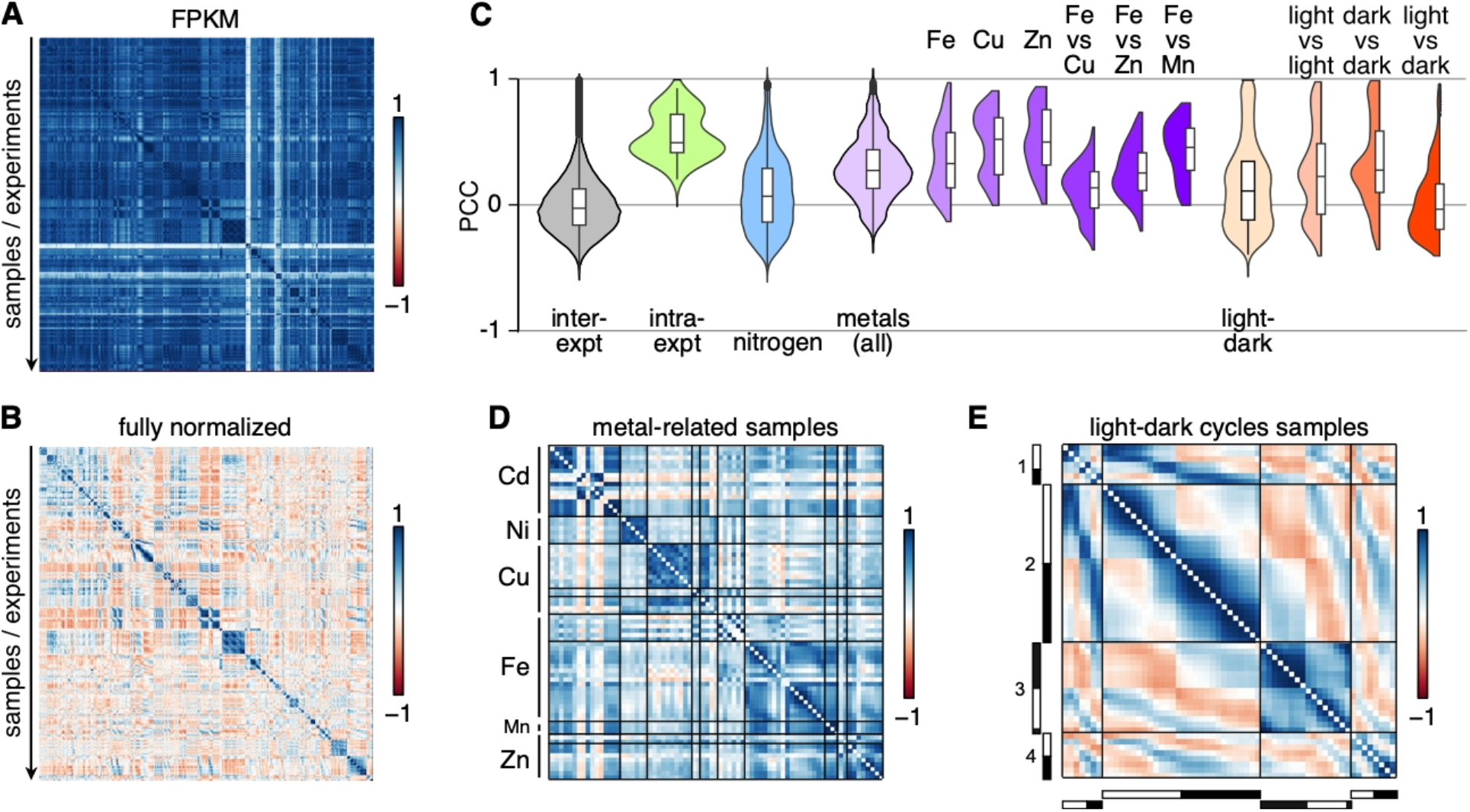
Samples from the Same Experiment are Strongly Correlated. Correlation matrices between all samples using expression estimates for all 17,741 nuclear genes as FPKM **(A)**, or after all normalization steps **(B)**. Samples belonging to the same experiment are in consecutive order, and roughly in chronological order. **(C)** Ditribution of Pearson’s correlation coefficients between (“inter-expt”) and within (“intra-expt”) experiments. PCCs for all comparisons between experiments are shown as violin plots and box plots (“inter-expt”, gray), alongside mean PCCs from all samples within each experiment (“intra-expt”, green), samples collected in the context of nitrogen deprivation (blue), PCCs for all metal-related samples (light purple) and specific metals (darker shades of purple), samples collected over a diurnal cycle (light orange) and PCC between subsets of samples (darker shades of orange). Values along the diagonal of the matrix (equal to 1) were discarded prior to plotting. **(D)** Correlation matrix for samples from metal-related experiments, all from the Merchant laboratory, and in which either one micronutrient has been omitted from the growth medium (for deficiency conditions: copper Cu, iron Fe, manganese Mn and zinc Zn) or a toxic metal was added to observe the effect on homeostasis (cadmium Cd and nickel Ni). **(E)** Correlation matrix of samples collected over a diurnal cycle. The light- and dark-part of each sampling day is indicated on the left and bottom sides of the matrix as white and black bars, respectively. Four time-courses are compared here (Zones et al., 2015; Strenkert et al., 2019b; Panchy et al., 2014).

Two sets of experiments deviated from the general trend: experiments that were 1) metal-related (Figure 1D) or 2) that spanned a diurnal cycle (Figure 1E). Positive correlations largely segregated samples collected from cultures lacking a single micronutrient (copper Cu, iron Fe, manganese Mn and zinc Zn) into their targeted deficiency. Based on correlations across samples, Fe-deficient cultures were slightly more similar to Zn- and Mn-deficient cultures than they were to Cu-deficient cultures (Figure 1C), as expected. These observations support the hypothesis that these three metals (Fe, Zn and Mn) are transported by partially overlapping sets of transporters and involve partially shared regulon components (Tsednee et al., 2019; Merchant et al., 2006; Malasarn et al., 2013; Hong-Hermesdorf et al., 2014a). Metal-related experiments appeared more related to each other than to any other experiments, which may reflect the tightly-controlled growth conditions we follow for such studies (Moseley et al., 2002a; Urzica et al., 2012b; Hong-Hermesdorf et al., 2014b; Allen et al., 2007). However, these correlations clearly did not extend to non-metal related work within our own laboratory, despite using the same stock solutions, growth medium recipes, and incubators (Supplemental Figure 4E, 4F).

The correlation matrix between diurnal samples was striking: we obtained the highest degree of positive correlation between samples that were temporally close to one another within and across diurnal experiments (Figure 2E). At a slightly broader scale, samples collected during the day were generally positively correlated, again within and across diurnal experiments, although the extent of correlation was stronger between samples from the same experiment. The same observation held true when comparing samples collected during the night part of the diurnal cycle. Finally, samples collected during the day were negatively correlated with samples collected at night, both within and across experiments (Figure 1E). In all diurnal samples, over 80% of nuclear genes exhibited a rhythmic pattern with phases spanning the entire day (Strenkert et al., 2019; Zones et al., 2015). That diurnal samples can cluster so clearly according to their collection time suggests that the endogenous timing of an unknown sample might be accessible by comparing its correlation profile with that of known diurnal datasets. This approach is similar in concept to the molecular timetable method used to detect sample time from single time-point data (Ueda et al., 2004).

**Figure 2.**
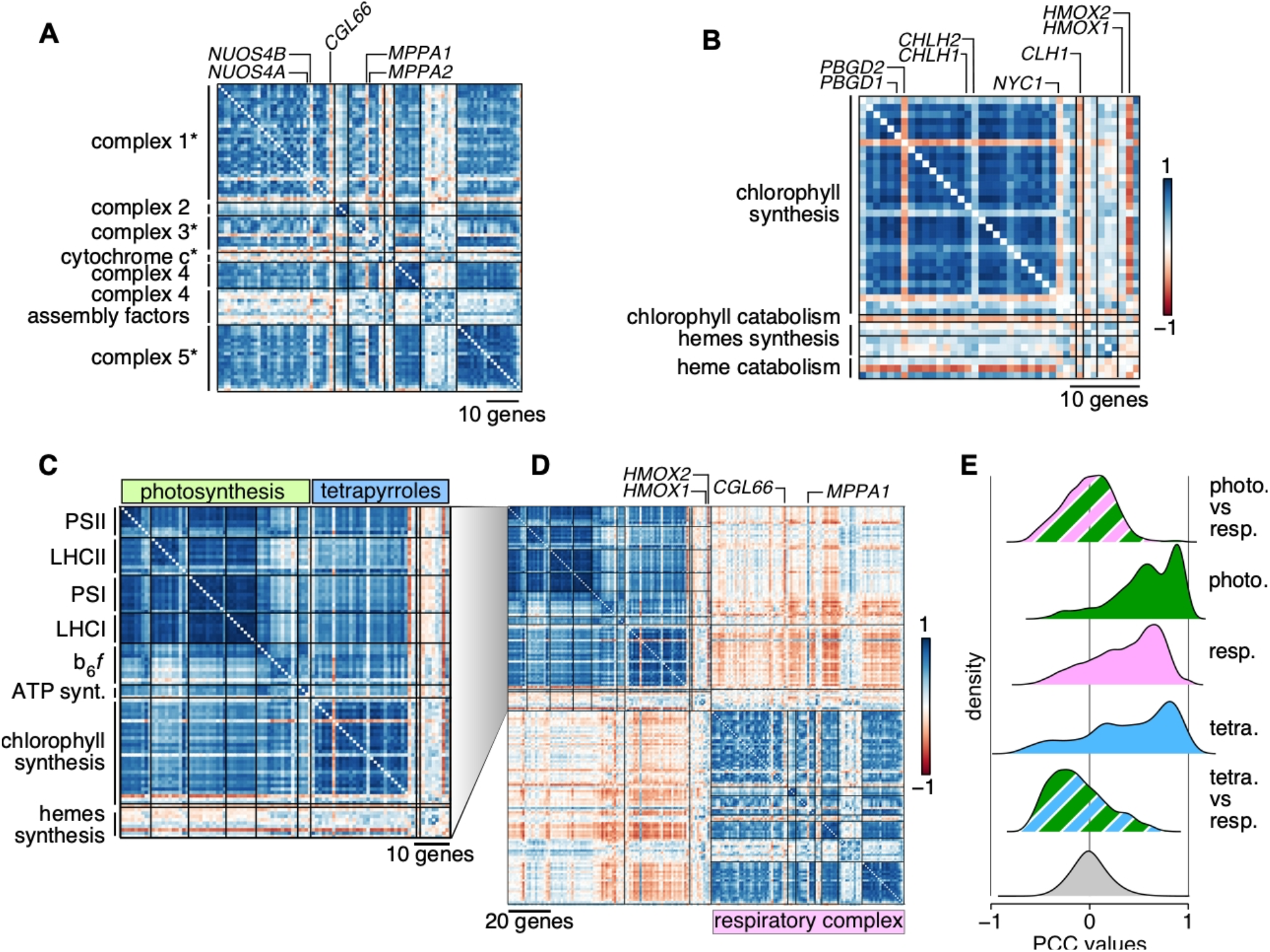
Correlations and Anti-Correlations between Organellar Energy Producing Systems. **(A)** Correlation matrix of nucleus-encoded components of each mitochondrial respiratory complexes, in the order defined by Zones et al. (Zones et al., 2015). An asterisk after the name of a complex signifies that its dedicated assembly factors (one to two genes outside of complex 4) are shown last, after the complex components. **(B)** Correlation matrix of chlorophyll and hemes biosynthetis genes. Genes have been ordered according to Zones et al., (2015). Pairs of homologous genes are indicated above the correlation matrix. **(C)** Co-expression matrix of photosystem genes (in green) and tetrapyrroles biosynthetic genes (in blue). **(D)** Comparison of co-expression profiles of chloroplast- and mitochrondrion-localized energy production systems. The respiratory complex matrix is redrawn from Supplemental Figure 9. **(E)** Distribution of PCCs between groups of genes. The gray distribution is the genome-wide distribution of all PCCs between all gene pairs. Photosynthesis: photo.; tetrapyrroles: tetra.; respiration: resp..

### Co-Expression Potential in Manually Curated Gene Lists

We next turned our attention to correlation between genes to dissect co-expression potential in Chlamydomonas. We calculated PCCs for all gene pairs (157,362,670 pairs, not counting self-self pairs); they followed a normal distribution (Kolmogorov-Smirnov test statistic D = 0.019, p-value < 2.2 × 10^−16^), indicating that most gene pairs are not co-expressed (Supplemental Figure 5A). Although the distribution of all PCC values had a mean of zero, its two tails contained the most interesting gene pairs with high absolute correlations. Of the 157,362,670 possible gene pairings, 5.4 % (or 845,249 gene pairs) had PCC values < –0.6 and > +0.6. Fewer gene pairs were defined by PCC values < –0.8 and > +0.8, accounting for 0.5 % of all PCCs (or 76,462 pairs), nevertheless leaving ample room for co-expression.

Hierarchical clustering suggested that sets of genes displayed very similar expression behaviors, as the larger blue blocks visible along the diagonal of the correlation matrix attested (Supplemental Figure 5B and 5C). Based on these observations, we followed a three-pronged approach to test for co-expression and identify co-expressed genes. First, we determined the extent of co-expression and anti-correlation in manually-curated gene lists from the community. Second, we defined the co-expression cohort associated with a given nuclear gene. Third, we identified co-expression modules. Both latter approaches entailed calculating the Mutual Rank (MR) associated with each gene pair (Obayashi and Kinoshita, 2009; Aoki et al., 2016; Wisecaver et al., 2017). We then turned MRs into edge weights as a measure of the connection between co-expressed genes (or nodes) for the construction of five MR-based co-expression networks with decreasing decay rates, denoted N1 to N5. During this process, we identified all genes that were co-expressed with each individual nuclear gene (Supplemental Data Sets 2-4 for networks N1-N3) and their anti-correlated cohorts, by inverting the rank order (Supplemental Data Sets 5-7). Each gene was at the center of a co-expression cohort with a clustering coefficient of zero (Supplemental Table 2). Faster decay rates restricted the size of co-expressed cohorts: with the most stringent criteria, a Chlamydomonas gene was co-expressed with 1 to 68 genes, with a mean cohort size of 17 genes. Relaxing the stringency imposed on co-expressed genes increased the mean size of cohorts to 36 (N2 networks) and 98 (N3 networks) (Supplemental Table 2).

As a proof of concept, we turned to gene lists compiled by the community. These lists comprised genes that participate in the same biological function or pathway, but information about their co-expression potential is incomplete. In particular, most co-expression analyses focus on positive correlations as the core criterion for the identification of co-expressed groups. Here, we capitalized on the graphical output of the R package *corrplot* to indicate 1) whether and 2) what fraction of genes was co-expressed, and 3) whether the expression profile of any gene within the lists was anti-correlated with others. We will acknowledge here that only genes with fairly dynamic expression profiles will register a co-expression pattern. By contrast, genes with low variance will have PCCs close to zero.

Since Chlamydomonas is a premier reference organism for organellar biogenesis, cilia biosynthesis and biology, we determined the co-expression potential of genes encoding components of the mitochondrial respiration chain, photosystems, chlorophyll and hemes biosynthesis (Figure 2), as well as motile cilia (Figure 3). We also assessed the co-expression potential of ribosome protein genes (*RPG*s) (Figure 4), as much early work in Chlamydomonas has described the organellar protein translation machinery in detail (Sager and Hamilton, 1967; Siersma and Chiang, 1971; Ohta et al., 1975; Martin et al., 1976). Finally, we tested co-expression between genes encoding transcription factors in Chlamydomonas and Arabidopsis (Figure 5).

**Figure 3.**
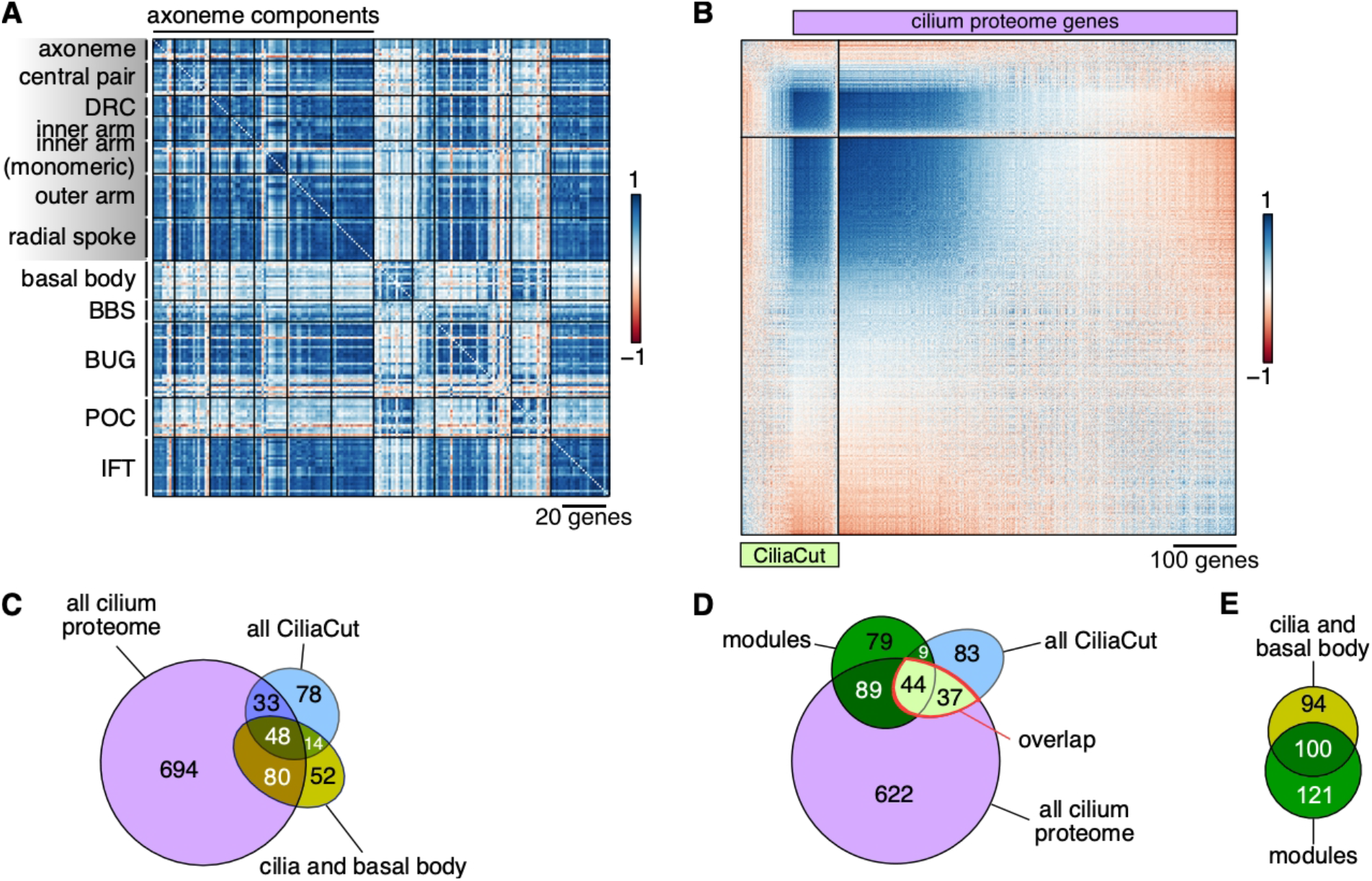
Confirmation of High-Confidence Cilium Proteins Based on Co-Expression of their Encoding Genes. **(A)** Correlation matrix of structural constituents of the Chlamydomonas cilia, in the order defined by Zones et al. (Zones et al., 2015). DRC: dynein regulatory complex; BBS: Bardet-Biedl syndrome protein complex; BUG: basal body upregulated after deflagellation; POC: proteome of centriole; IFT: intra-flagellar transport. **(B)** Correlation matrix between genes belonging to CiliaCut (green) or encoding components identified in the cilium proteome (light purple; Pazour et al., 2005). The genes within each subset were subjected to hierarchical clustering (First Principle Component (FPC) method in *corrplot*). **(C)** Venn diagram of the overlap between genes encoding putative components of the cilium proteome, CiliaCut and the cilia and basal body. Note that the gene lists do not reflect co-expression here. **(D)** Venn diagram of the overlap between genes encoding putative components of the cilium proteome, CiliaCut and genes belonging to cilia-related co-expression modules (listed in Suplemental Table 3). **(E)** Venn diagram of the overlap between genes encoding putative components of the the cilia and basal body and genes belonging to cilia-related co-expression modules.

**Figure 4.**
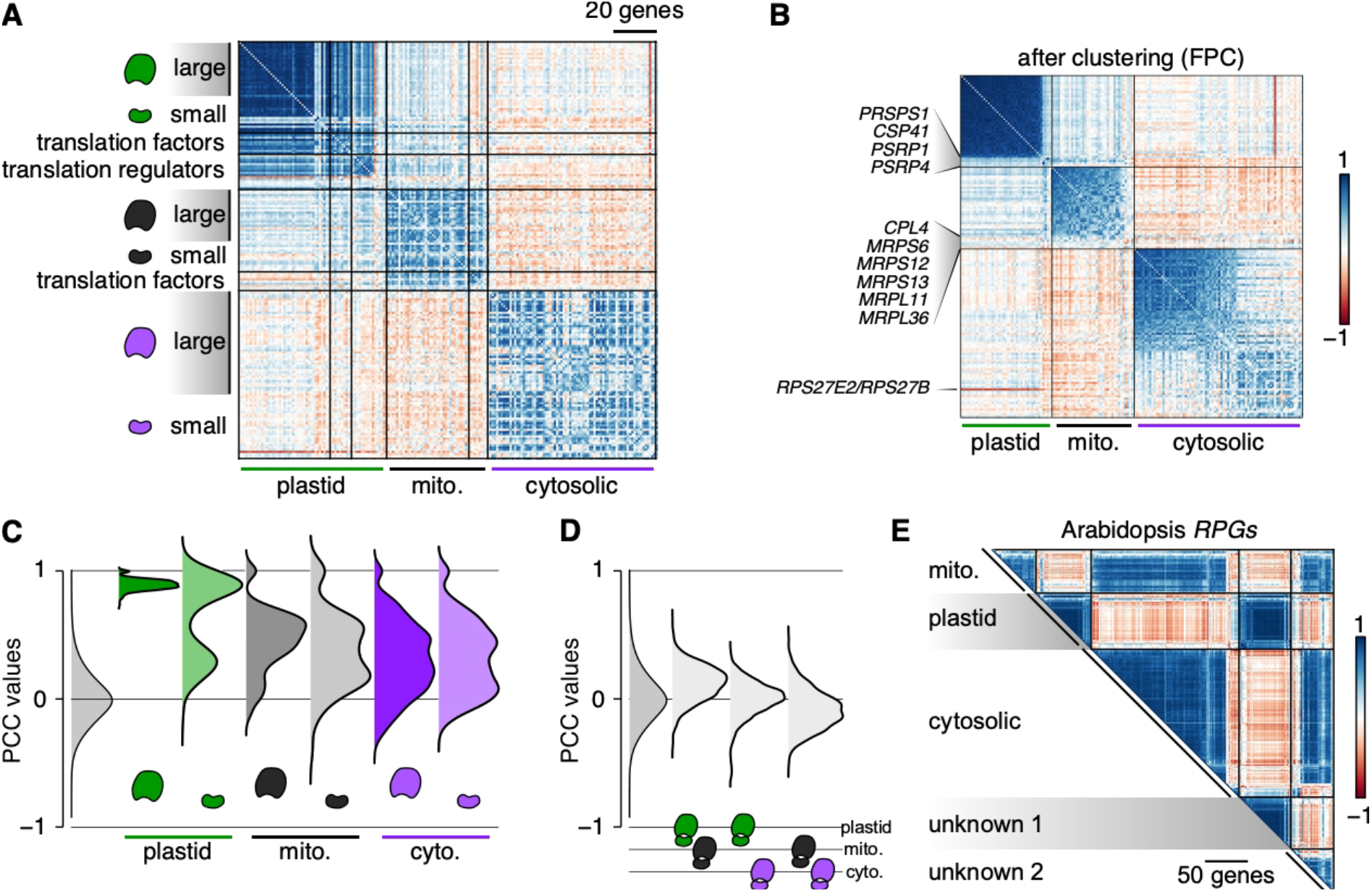
Co-Expression Between Ribosomal Protein Genes Reflects the Final Location of the Corresponding Ribosomal Proteins. **(A)** Correlation matrix between ribosomal protein genes (*RPGs*) and their translation regulators, sorted by the subcellular localization of their encoded proteins. For each set of *RPGs* and their regulators, we followed the same gene order defined by Zones et al. (Zones et al., 2015). **(B)** Correlation matrix restricted to *RPGs*. Each set of *RPGs* was subjected to hierarchical clustering (FPC method in *corrplot*) to single out non co-expressed genes. **(C)** Distribution of PCCs between *RPG* gene pairs encoding large or small ribosome subunits. The gray distribution indicates the PCC distribution of all gene pairs for the Chlamydomonas genome. **(D)** Distribution of PCCs for gene pairs belonging to distinct *RPG* groups. **(E)** Correlation matrix for 429 *RPGs* using the fully normalized dataset derived from Arabidopssi microarray experiments (Supplemental Data Set 7). “unknown 1” and “unknown 2” denote predicted *RPGs* whose encoded proteins have not been clearly assigned a localization. Note how “unknown 1” *RPGs* show strong correlation with chloroplast *RPGs* (cp), while “unknown 1” *RPGs* appear to be strongly correlated with cytosolic *RPGs*.

**Figure 5.**
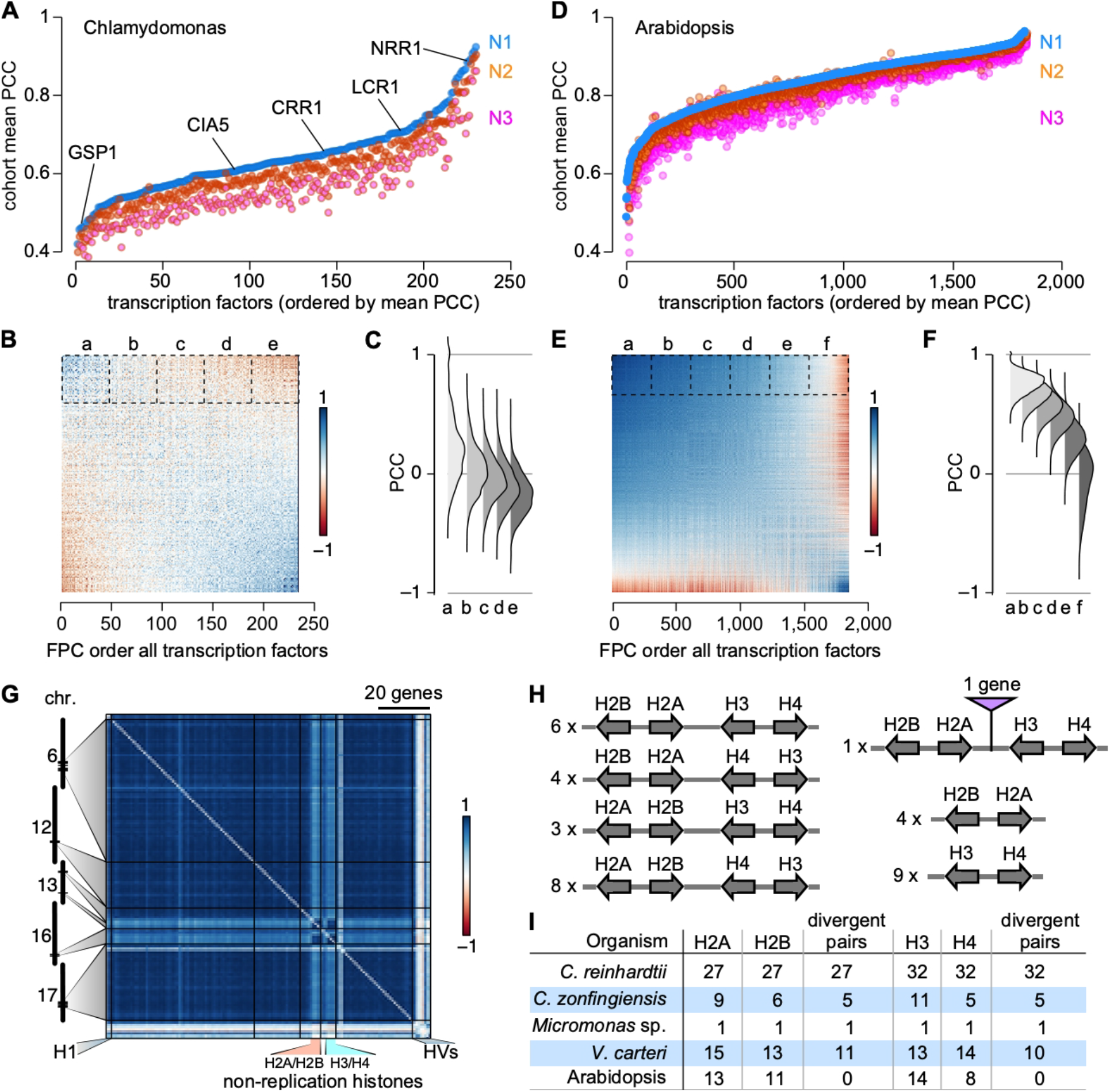
Correlations Between and Across Transcription Factors in Chlamydomonas and Arabidopsis, and the Special Case of Chlamydomonas Histone Genes. **(A)** Ordered mean Pearson’s correlation coefficient (PCC) for each Chlamydomonas gene encoding a DNA-binding protein or a transcription factor. We calculated the mean correlation between each gene and their co-expressed cohort (from networks N1, N2 or N3, as indicated). PCCs from networks N2 and N3 were ordered according to increasing mean PCC from network N1 co-expressed cohorts. Several transcription factors are listed for reference. **(B)** Correlation matrix between Chlamydomonas genes encoding a DNA-binding protein or a transcription factor, ordered according to First Principle Component (FPC) clustering method built in *corrplot*. **(C)** Distribution of inter-transcription factor PCCs plotted in (**B**). We defined five groups, indicating by a-e in (**B**) and (**C**). **(D)** Ordered mean Pearson’s correlation coefficient (PCC) for Arabidopsis genes encoding a DNA-binding protein or a transcription factor with a probe on the AtH1 Affymetrix microarray. Genes were ordered based on increasing PCC from network N1 co-expressed cohorts. **(E)** Correlation matrix between Arabidopsis genes encoding a DNA-binding protein or a transcription factor, ordered according to the FPC clustering method built in *corrplot*. **(F)** Distribution of inter-transcription factor PCCs plotted in (**E**). We defined six groups, indicating by a-f in (**E**) and (**F**). **(G)** Correlation matrix among Chlamydomonas histone genes, ordered according to their genomic coordinates. Histone genes that are not regulated by the cell cycle are indicated as “non-replication histones”. **(H)** Global clustering of histone genes in Chlamydomonas. All histone genes occur as divergent pairs, and are oftentimes grouped as one representative of each major histone type (H2A, H2B, H3 and H4). The number to the left gives the number of instances of the given arrangement in the Chlamydomonas genome. **(I)** Comparison of histone gene clustering in selected photosynthetic organisms. *V. carteri*: *Volvox carteri*; *C. zofingiensis*: *Chromochloris zofingiensis*.

#### Nucleus-encoded organellar energy systems

Mitochondria and chloroplasts provide energy and reducing power to the cell, albeit at different times of day. Based on previous results (Strenkert et al., 2019; Zones et al., 2015), we expected to observe global co-expression of genes encoding components of the respiratory complex. Indeed, most genes whose products participate in electron transport or oxidative phosphorylation were co-expressed (Figure 2A), although some genes deviated from this pattern. For instance, *CONSERVED IN THE GREEN LINEAGE 66* (*CGL66*, Cre09.g390467) was negatively correlated with other complex 1 genes, suggesting that it may not belong to this complex, or functions as a negative regulator. Proteins encoded by two related genes provided an example of potential sub-functionalization: *NUOS4B* (Cre16.g681700, from complex 1) and *MITOCHONDRIAL PROCESSING PEPTIDASE ALPHA SUBUNIT* (*MPPA1*, Cre17.g722800, from complex 3) were not co-expressed with other genes coding for components forming their respective complexes, although the related genes *NUOS4A* and *MPPA2* were (and were also more highly expressed).

Of the genes involved in tetrapyrroles biosynthesis, only those encoding enzymes responsible for chlorophyll biosynthesis appeared to be co-expressed, with the exception of the porphobilinogen deaminase gene *PBGD2* (Cre02.g113850) and the magnesium chelatase subunit H gene *CHLH2* (Cre11.g4776625), although their homologues *PBGD1* and *CHLH1* were (Figure 2B), with *PBGD1* expressed at much higher levels than *PBGD2*. By contrast, heme biosynthetic genes exhibited no co-expression with genes from either photosystem (mean PCC: −0.03 ± 0.23).

All photosynthetic genes were strongly co-expressed (Figure 2B). Although heme and chlorophyll biosynthesis compete for the same pool of precursors, the expression of the genes involved in each pathway is independent (mean PCC: 0.04 ± 0.28). Genes encoding heme-containing enzymes and other cytochromes were however anti-correlated with chlorophyll biosynthetic genes (Figure 2B-2D), thereby ensuring that adequate levels of heme be synthesized without reaching toxic levels by coordinating the heme pool with heme binding proteins. The two heme oxygenase genes followed distinct expression behaviors: *HMOX1* was weakly co-expressed with photosystems and other tetrapyrrole biosynthetic genes, whereas *HMOX2* was strongly anti-correlated with them, consistent with the light-dependent repression of this gene (Wittkopp et al., 2017). Furthermore, the *hmox1* mutant is pale-green, a phenotype typical for chlorophyll biosynthesis mutants. Notably, the expression of genes involved in photosynthesis is not affected in the *hmox1* background, which is consistent with the general lack of correlation between *HMOX1* and photosystems (Wittkopp et al., 2017).

Finally, genes encoding proteins that form the mitochondrial respiratory complex were largely anti-correlated with photosynthetic and tetrapyrrole biosynthetic genes (Figure 2D, 2E). This opposite co-expression may partially stem from the distinct temporal separation of the underlying cellular events: high expression during the day for photosynthesis and tetrapyrroles biosynthesis, and high expression at night for mitochondrial respiration (Strenkert et al., 2019; Zones et al., 2015).

#### Cilia

The components of the Chlamydomonas cilia are coordinately transcribed following cell division at night, as cells first resorb their existing flagella prior to division and must synthesize them afresh in anticipation of dawn and photosynthetic activity (Cross and Umen, 2015; Wood et al., 2012; Rosenbaum et al., 1969). Although most RNAseq samples were collected from cultures grown in constant light and, presumably, asynchronous, we observed strong co-expression across most genes encoding structural components of the cilia (mean PCC: 0.65 ± 0.18), as well as with components of IntraFlagellar Transport (IFT) particles responsible for the assembly, maintenance and signaling within cilia (mean PCC: 0.74 ± 0.17) (Figure 3A). Several genes did not follow this general trend: they encoded proteins that modify protein function and therefore act at the post-translational level (Flagella Associated Protein 8 (FAP8), a protein phosphatase 2A regulator; enolase, contributing to ATP production within cilia, and a number of chaperones or heat shock proteins [DNJ1, HSP70A]). Other genes that were not co-expressed encoded proteins with cellular roles outside of cilia, for instance HSP70A, actin and profilin, suggesting that a fraction of the total pool of each protein participates in cilia biogenesis while the bulk carries out functions in the cytosol.

Centriole proteins have been identified by a number of techniques, including mass spectrometry of purified centrioles, co-expression following deflagellation, and comparative genomics (Keller et al., 2005; Keller and Marshall, 2008; Li et al., 2004). Genes encoding most basal body components were indeed co-expressed across all our samples and showed strong co-expression with *PROTEOME OF CENTRIOLE* (*POC*) genes. Both basal body and *POC* genes were however only weakly co-expressed with genes coding for cilia components, as might be expected: the centriole is always present in the cell, whereas cilia form a more dynamic structure (Figure 3A). As previously described, the majority of *BASAL BODY UPREGULATED AFTER DEFLAGELLATION* (*BUG*) genes were more co-expressed with cilia components than with basal body markers (Figure 3A). The co-expression profile of several *BUG* genes (*BUG23*, *BUG24*, *BUG27*) suggested that their function may be instead associated with the centriole proper, as they showed stronger co-expression with basal body genes. We also denoted a lack of co-expression between basal body components and *CCT3*, *HSP90A*, *FMO11* and *PHB1*, all predicted to perform function(s) outside of the centriole (Zones et al., 2015).

Genes encoding components of the Bardet-Biedl syndrome protein complex (BBSome) were only weakly co-expressed (mean PCC: 0.29 ± 0.16) and were not co-expressed with basal body constituents (mean PCC: 0.23 ± 0.16), while moderately with ciliary structures (mean PCC: 0.38 ± 0.23). Our co-expression analysis of cilia and centriole components therefore accurately grouped genes based on function and cellular localization and highlighted those genes with distinct expression profiles. The ability to identify bona fide cilia and centriole components based on co-expression also offered the opportunity to subject larger lists to a similar analysis. The cilium proteome is predicted to comprise close to a thousand proteins based on proteomics analysis (Pazour et al., 2005), although a fraction is likely to correspond to contaminants. Likewise, a comparative genomics approach uncovered around 200 genes conserved between ciliated species and absent in all other species: Ciliacut (Li et al., 2004). These two lists overlap only partially, with 81 genes belonging to both. We wondered if co-expression profiling might allow to pull high-confidence cilia components: we measured co-expression in three groups (Ciliacut only; Ciliacut+cilium proteome overlap; cilium proteome only). The resulting correlation matrix is shown in Figure 4B. Genes only included in Ciliacut were on average not co-expressed with each other (mean PCC: 0. 03 ± 0.24) and consisted of many *MOTILITY* (*MOT*) genes not found in Caenorhabditis elegans (which lacks motile cilia) and *SENSORY, STRUCTURAL AND ASSEMBLY* (*SSA*) genes. Similarly, about 550 genes only present in the cilium proteome gene list showed no pattern of co-expression, with a mean PCC of 0.01 ± 0.22. In sharp contrast, 76 genes that belonged to both lists were highly co-expressed (mean PCC: 0.63 ± 0.20). Equally highly co-expressed was a set of ~300 genes whose encoded proteins are only found in the cilium proteome (mean PCC: 0.63 ± 0.15), with many uncharacterized *FLAGELLAR ASSOCIATED PROTEIN* (*FAP*) genes. Together, these two sets comprised over 400 co-expressed genes that are prime candidates for functional dissection. They are listed in Supplemental Data Set 8.

#### Ribosome Protein Genes

Nucleus-encoded ribosomal protein genes (*RPG*s) code for proteins with three cellular destinations. The co-expression pattern observed between *RPG*s largely reflected the organelle in which their encoded subunits will function (Figure 4A). Plastid *RPG*s exhibited the strongest degree of co-expression (mean PCC = 0.88 ± 0.06). The sole exceptions were *PLASTID SPECIFIC RIBOSOMAL PROTEIN1 PSRP1* and *PSRP4*, which are among the lowest expressed genes encoding small subunits proteins, and the gene encoding the Chloroplast Stem-loop binding Protein of 41 kDa CSP41 (mean PCC = 0.27 ± 0.09) (Figure 4B). Neither PSRP1 or CSP41 are thought to be plastid ribosomal proteins, but both participate in efficient translation, either by inducing conformational changes within the ribosome (PSRP1, Sharma et al., 2010) or by stabilizing target plastid RNAs (CSP41, Qi et al., 2012). Large and small plastid ribosomal subunits were co-expressed equally strongly (*PRPL*s: 0.89 ± 0.04; *PRPS*s: 0.86 ± 0.09 excluding *PSRP1* and *PSRP4*) (Figure 4C). Plastid translation factors also displayed a high degree of co-expression with one another (mean PCC: 0.52 ± 0.18) and with plastid *RPG*s (mean PCC: 0.59 ± 0.20). co-expression between chloroplast translation regulators defined three sub-groups: one group that was highly co-expressed with plastid *RPG*s (11 genes), one group that was not co-expressed (4 genes: *RNA-BINDING PROTEIN 38 RB38*, *ACETATE REQUIRING 115 AC115*, *BUNDLE SHEATH DEFECTIVE2 BSD2* and *CHLOROPLAST RHODANESE-LIKE TRANSLATION CRLT*), and a single weakly anti-correlated gene with all plastid *RPG*s, the translation factor and translation regulator (*TBA1*; mean PCC against *RPG*s: −0.35 ± 0.19).

The co-expression of *RPG*s encoding proteins destined for the mitochondrion or cytosol was less pronounced, but similar between large and small subunits *RPG*s (Figure 4C). For both compartments, correlation coefficients between *RPG*s followed a bimodal distribution, with a fraction of PCCs around zero. For mitochondrial *RPG*s, high expression levels appeared to come at the cost of lower PCCs, whereas the opposite was true for cytosolic *RPG*s. Mitochondrial *RPG*s tended to be weakly co-expressed with plastid *RPG*s (mean PCC: 0.13 ± 0.14) while anti-correlated with cytosolic *RPG*s (mean PCC: −0.08 ± 0.15) (Figure 4D). There was no clear correlation between the expression of most plastid and cytosolic *RPG*s (mean PCC: −0.0006 ± 0.14) (Figure 4D). As the single exception, the cytosolic *RPG RPS27E2/RPS27B*, which is generally expressed at much lower levels than all other cytosolic *RPG*s, stood out with a pronounced anti-correlation with plastid *RPG*s (mean PCC: −0.54 ± 0.05) (Figure 4B). Nitrogen deficiency results in a sharp increase in *RPS27E2* expression, concomitant with a global arrest in plastid translation until more auspicious conditions return (Schmollinger et al., 2014; Kajikawa et al., 2015; Plumley and Schmidt, 1989), which may explain the pattern observed here.

Given the strong correlation between sets of *RPG*s in Chlamydomonas, we wondered how conserved this pattern might be. We determined the correlation between Arabidopsis *RPG*s using normalized data from microarrays downloaded from AtGenExpress. The Arabidopsis genome contains 429 *RPG*s; as in Chlamydomonas, their encoded products will locate to one of three compartments (cytosol, mitochondria or chloroplasts). We observed a correlation matrix very reminiscent of that of Chlamydomonas *RPG*s: indeed, each organellar *RPG* set is co-expressed. A subset of Arabidopsis *RPG*s lacked a clear functional localization; however, co-expression with other *RPG*s clearly predicted their localization as being either plastidic (“unknown 1”) or cytosolic (“unknown 2” in Figure 4E). We provide the Arabidopsis normalized microarray dataset as Supplemental Data Set 9. We obtained similar results with *Physcomitrium patens* (not shown), although the exact interpretation is likely muddled by the multiple splice variants listed for each gene.

#### Transcription factors

As regulators of gene expression, transcription factors and other DNA-binding proteins will bind to their cognate cis-regulatory elements to modulate gene expression. We wished to test whether co-expression cohorts associated with transcription factors may help in deciphering their biological function. To this end, we calculated the mean PCC between a given transcription factor and its co-expressed cohort from networks N1, N2 and N3. As shown in Figure 5A, PCC values ranged from 0.42 to 0.92, with a mean of 0.64 ± 0.09. The gene encoding the transcription factor NITROGEN RESPONSIVE REGULATOR (NRR1) showed one of the highest PCCs (0.885 for its N1 cohort) and was highly co-expressed with two other transcription factor genes, both encoding Helix-Loop-Helix proteins (Cre01.g011150 and Cre04.g216200. The genes *LOW-CO2 RESPONSE REGULATOR* (*LCR1*) and *CIA5/CCM1* participate in the induction of gene expression in response to low CO_2_ conditions (Fang et al., 2012; Xiang et al., 2001; Fukuzawa et al., 2001; Yoshioka et al., 2004), with LCR1 predicted to act downstream of CIA5 (Yoshioka et al., 2004). Both genes showed high correlation with their N1 cohorts (*CIA5*: mean PCC of 0.61 with 20 genes and *LCR1*: mean PCC of 0.715 with 34 genes), but their cohorts did not overlap. In addition, we failed to identify *LCR1* as a gene co-expressed with *CIA5*, suggesting that each transcription factor may act in parallel rather than in converging pathways. We also looked at the extent of co-expression between transcription factors, as illustrated in Figure 5B and 5C. When subjected to hierarchical clustering with the First Principle Component (FPC) method from *corrplot*, transcription factor genes showed weak to moderate co-expression, as well as anti-correlations. The potential for co-expression (or anti-correlation) did not appear to follow simple rules related to the family of the transcription factors. The dataset generated here will provide an interesting opportunity to compare the output from methods such as DNA Affinity Purification and sequencing (DAP-Seq) (O’Malley et al., 2016).

We performed the same analysis with Arabidopsis transcription factors. We calculated the mean PCC for 1,864 transcription factors represented by a probe on the ATH1 Affymetrix microarray: mean PCCs per gene ranged from 0.49 to 0.96, with a mean of 0.86 ± 0.06 (Figure 5D. Co-expression between Arabidopsis transcription factors was much more evident than in Arabidopsis, as shown in Figure 5E and 5F. The vast majority of genes encoding transcription factors showed strong co-expression, but a subset was anti-correlated (Figure 5E) that was not explained based on a crude separation into leaf- and root-specific expression pattern. We thus selected a list of 150 genes exhibiting the highest anti-correlation and subjected them to Gene Ontology (GO) enrichment analysis to determine their function. The biological processes associated with these genes included “heterochronic regulation of development”, “photoperiodism”, “regulation of seed development” and “regulation of flower development”, raising the possibility that the observed pattern may reflect the temporal rather than the spatial specificity of regulatory proteins.

Returning to Chlamydomonas genes encoding DNA-binding proteins, we took a closer look as histone genes, most of which are coordinately expressed with a peak in expression shortly before cell division (Strenkert et al., 2019a; Zones et al., 2015). However, a small group of histone genes remain constantly expressed over the diurnal cycle and are termed “non-replication” (or emergency) histones. Histone genes displayed a striking co-expression pattern, with all replication histones being highly co-expressed (Figure 5G). Similarly, non-replication histones were strongly co-expressed as a group, but less so when probed against replication histones. Histone variants showed little correlation in their expression with either group. While assembling the gene list for histones, we noticed their high numbers (117 histone genes, not counting histone variants) and their tight clustering along only five chromosomes (Figure 5G). Even more remarkable was their arrangement as divergent gene pairs: all histone H2A and H2B genes were present as divergent pairs, and all histone H3 genes occurred as a divergent partner to a histone H4 gene. In many cases, each major histone class was represented in a 4-gene cluster, corresponding to 84 (out of 117) histone genes (Figure 5H). The high number of histone genes appeared to be unique to Chlamydomonas, as the genomes of the other unicellular algae *Micromonas* sp., *Chromochloris zofingiensis* and *Volvox carteri* encoded far fewer histones. However, the clustering of histones as divergent gene pairs was partially maintained, as summarized in Figure 5I. In *Micromonas*, the four histone genes were arranged as two divergent pairs, with H2A and H2B belonging to one pair, and H3 and H4 found in the second pair. Likewise, most histone genes from *C*. *zofingiensis* and *V. carteri* grouped in divergent pairs.

### Co-Expression Cohorts and Co-Expression Modules

Testing co-expression between members of a gene family may help assign specific functions, or group them in functionally homologous groups. We applied the co-expression cohort approach to the ferredoxin gene family, consisting of 12 genes (*FDX1-FDX12*). FDX1, also known as PETF, is the main electron acceptor from Photosystem I during photosynthesis. FDX5 has been shown to function in fatty acid desaturation (Yang et al., 2015), while FDX9 likely plays a critical role in fermentation (Strenkert et al., 2019b). We extracted the co-expression cohort associated with each *FDX* from network N1 (Supplemental Data Set 10), and plotted the complete correlation matrix, as shown in Supplemental Figure 6. As expected, the *FDX1* cohort included the FDX1 partner ferredoxin *NADP reductase* (*FNR1*) and genes encoding multiple subunits of cytochrome b_*6*_*f*. Each *FDX* cohort varied in size and in the function of its constituents. For instance, *FDX4* showed strong co-expression with several tetrapyrroles biosynthetic genes, while *FDX2* was co-expressed with the nitrite transporter *NAR1.6* and the nitrate transporter *NAR2*. In addition, *FDX1* and *FDX2* were anti-correlated (PCC: −0.48), as were their respective cohorts (Supplemental Figure 6), pointing to specific functions for FDX2 outside of photosynthesis. *FDX5* was shown previously to be induced under Cu deficiency, and its co-expression cohort comprised several genes up-regulated in low-Cu conditions, including the putative Cu transporters *CTR1* and *CTR2* (Page et al., 2009), as well as *COPPER RESPONSE DEFECT1* (*CRD1*), which encodes a chlorophyll biosynthetic gene that functions specifically under low-Cu conditions (Moseley et al., 2002b). *FDX6* was itself co-expressed with several genes involved in carotenoid biosynthesis, suggesting a role for the protein.

We conclude that co-expression cohorts can provide useful information when characterizing a gene of interest, and may offer hints about the underlying function of the encoded protein. For example, we validated a role for FDX2 in nitrogen metabolism based on co-expression alone, corroborating earlier results (Terauchi et al., 2009). We also suggest that FDX4 may participate in tetrapyrroles biosynthesis.

We next used our co-expression cohorts and associated edge weights as input for the graph-clustering Cytoscape plugin ClusterONE (Nepusz et al., 2012), resulting in the identification of 616 co-expressed modules for network N1, 248 modules for network N2, and 117 modules for network N3 (Supplemental Figure 7, Supplemental Table 2). We restricted our efforts to the N3 network as a good compromise between larger module sizes and significant GO enrichment within modules. Out of 117 N3 modules, we grouped 37 modules into 8 functional groups based on their significant enrichment in biological processes: transcription, translation, ribosome biogenesis, protein degradation, DNA replication, transport, photosynthesis and flagella biogenesis and function (Supplemental Table 3, Supplemental Data Set 11). A single module defined a ninth group associated with response to phytohormones, specifically cytokinin, whose signaling cascade is incomplete in the microalga (Lu and Xu, 2015). These categories were not surprising and satisfying all the same: they broadly mapped to conserved cellular functions, or to processes where Chlamydomonas is a premier model organism for their study.

To obtain genes that are co-expressed with a list of interest, we separately used manually-curated gene lists as baits to extract their co-expressed genes from the N1, N2 and N3 networks. As stringency decreases from the N1 to the N3 networks, the number of selected genes increased, but the resulting lists were nested. Co-expression cohorts associated with gene lists expanded the number of potentially informative genes 2-20 fold, with an average increase of 10-fold (Supplemental Figure 8). Using genes from co-expression modules as baits, we thus identified their associated co-expressed cohorts and determined the extent of overlap with other user-defined lists (as illustrated in Figure 3C) to obtain high-confidence genes. We also established the timing of peak expression over the diurnal cycle for each module, group and co-expressed cohorts, using the diurnal phase of all genes considered rhythmic in two diurnal datasets (Supplemental Figure 9) (Zones et al., 2015, Strenkert et al., 2019).

#### Cell Division Modules

Five modules involved in cell division and DNA replication comprised a non-redundant set of 245 genes (Figure 6A), with 88 genes with an acronym and 157 with no prior functional knowledge. Using guilt by association, we propose that these non-annotated genes play a role in some aspect of cell division. Only 19 out of the 245 genes overlapped with 79 genes identified by forward genetic screens for defects in cell cycle progression; this overlap was limited to the highly co-expressed genes within both sets (Figure 4A) (Tulin and Cross, 2014; Breker et al., 2018). We then determined the co-expression cohorts associated with each gene list and assessed their overlap. By definition, all genes within our modules are highly inter-connected, but they also exhibited co-expression with ~ 400 additional genes that define a larger cohort with presumptive function in cell division (Figure 6B). Similarly, hundreds of genes showed strong co-expression with the 30 co-expressed genes from the genetics list (Figure 6C). Finally, we defined a third list comprising genes critical for DNA replication, chromosome segregation and cell division proper, for which we determined the co-expression cohorts (Figure 6D, Supplemental Data Set 12).Notably, although the initial gene lists were quite distinct (Figure 6E), their cohorts shared more genes as network stringency decreased, suggesting that the intersection of co-expression cohorts converged on a common set of genes.

**Figure 6.**
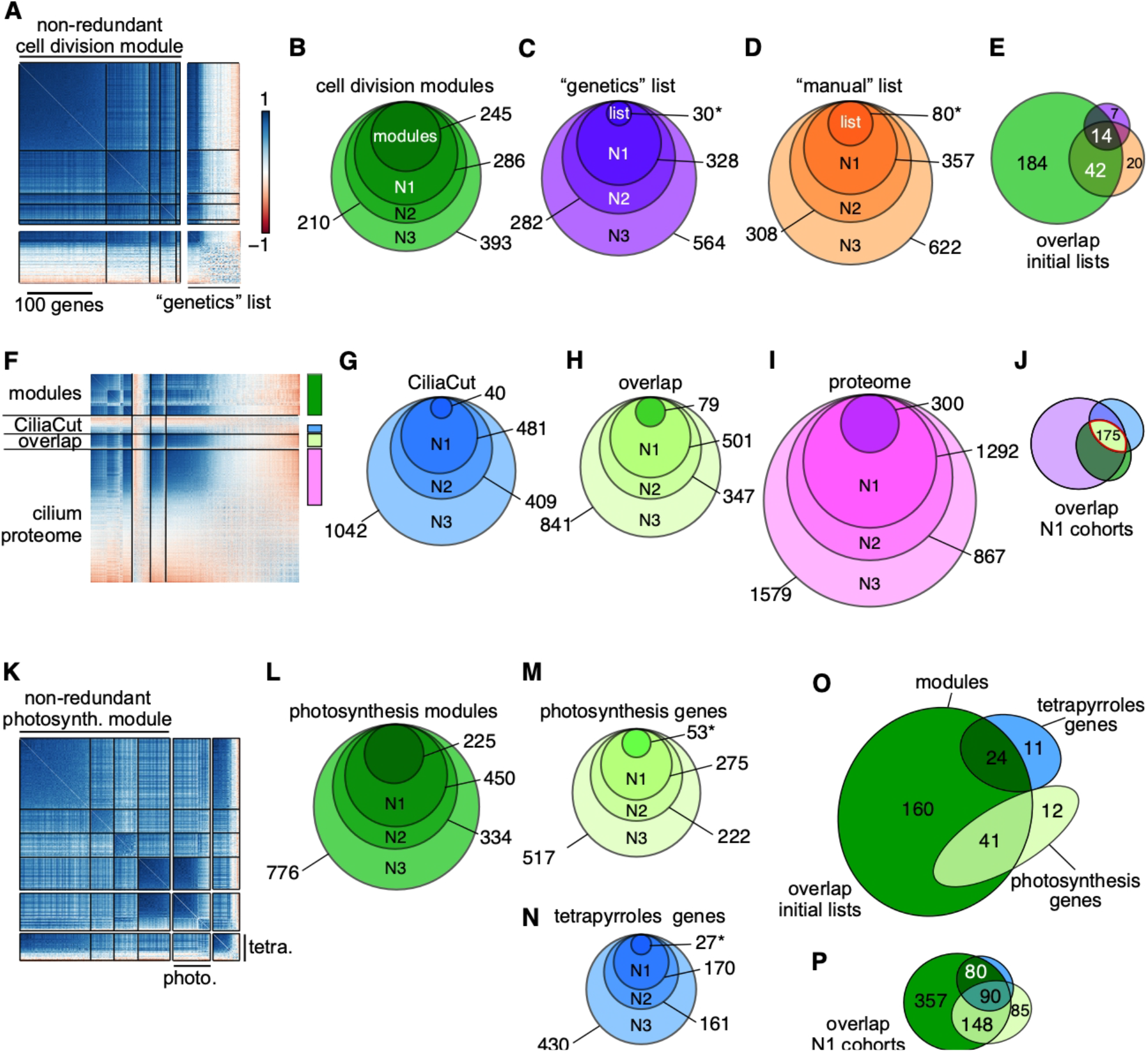
Core cell division genes are coordinately and highly co-expressed. **(A)** Correlation matrix of non-redundant cell division modules and correlation matrix of genes whose loss of function leads to cell division defects (Tulin and Cross, 2014; Breker et al., 2018). Genes within each set were ordered according to hierarchical clustering using the FPC method in *corrplot*. **(B-D)** Co-expressed cohorts, shown as nested Venn diagrams, associated with genes from the cell division modules **(B)**, the genetics list **(C)** or genes involved in DNA replication and chromosome segregation (manual list) **(D)** from networks N1-N3. **(E)** Overlap between original gene lists related to cell division (modules, genetics and manual lists). **(F)** Correlation matrix of non-redundant cilia modules (“modules”) and genes belonging to CiliaCut only (“CiliaCut”), the cilium proteome and shared genes between CiliaCut and the cilium proteome (“overlap”). The color bars on the right refer to the color scheme used for co-expression cohorts in **G-J**. **(G-I)** Co-expressed cohorts, shown as nested Venn diagrams, associated with genes from CiliaCut **(G)**, the overlap between CiliaCut and the cilium proteome **(H)** and the cilium proteome **(I)** from networks N1-N3. **(J)** Overlap between N1 cohorts associated with each initial gene list (CiliaCut, overlap and cilium proteome). **(K)** Correlation matrix of non-redundant photosynthesis modules, photosynthesis-related genes and tetrapyrrole biosynthesis-related genes. **(L-N)** Co-expressed cohorts, shown as nested Venn diagrams, associated with genes from the photosynthesis modules **(L)**, photosynthesis-related genes **(M)** and tetrapyrrole biosynthesis-related genes **(N)** from networks N1-N3. **(O)** Overlap between initial gene lists. **(P)** Overlap between N1 cohorts associated with photosynthesis and tetrapyrrole biosynthesis. In panels **C**, **D**, **M and N**, the asterisk indicates that the gene list was restricted to highly co-expressed genes, based on FPC clustering of the data.

#### Proteasome-Dependent Protein Degradation

Two modules shared a function in protein degradation. They largely overlapped and defined a set of 96 genes that included all but two of the 26S proteasome subunit genes. Most subunits of the 26S proteasome were highly co-expressed (mean PCC: 0.67 + 0.13). *CSN2* and *CSN6* were however not part of the protein degradation modules; they exhibited the weakest co-expression profile with other 26S proteasome subunit genes, although clearly still quite high (*CSN2* mean PCC: 0.54 ± 0.15; *CSN6* mean PCC: 0.53 ± 0.06) (Supplemental Figure 10A). The Chlamydomonas ortholog for the E3 ubiquitin ligase CONSTITUTIVE PHOTOMORPHOGENIC 1 (COP1), Cre13.g602700 (currently annotated as SPA1, Gabilly et al., 2019), showed no co-expression with the 26S proteasome (mean PCC: – 0.09 ± 0.10), consistent with a role as a regulatory component of the proteasome. We observed the same absence of co-expression in Arabidopsis between *COP1* and the remaining subunits of the proteasome, indicating a conserved mode of control from unicellular algae to land plants.

Proteasome-dependent proteolytic degradation entails the addition of ubiquitin onto the protein targeted for removal by the concerted action of E1 ubiquitin-activating enzymes, E2 ubiquitin-conjugating enzymes and E3 ubiquitin ligases. The Chlamydomonas genome bears 13 ubiquitin genes, three genes encoding potential E1 enzymes (Cre09.g386400, Cre06.g296983, and Cre12.g491500) and 17 genes coding for E2 enzymes. We did not compile a list of all E3 ubiquitin ligase genes, as they form large gene families. Our protein degradation modules only incorporated a single gene each for ubiquitin (*UBQ2*), E1 activating enzyme (Cre12.g491500, annotated as *UBA2*) and E2 conjugating enzyme (*UBC21*, although it was the second lowest-expressed *UBC* gene in our dataset; Supplemental Figure 10A). No other ubiquitin gene displayed a co-expression pattern with our protein degradation modules. By contrast, both remaining E1 enzyme genes (Cre09.g386400 and Cre06.g296983) were highly co-expressed with genes from our protein degradation modules. Likewise, we identified a subset of genes encoding E2 conjugating enzymes that were co-expressed with 26S proteasome subunit genes: *UBC3* (Cre03.g167000), *UBC9* (Cre16.g693700, also the most highly expressed *UBC* gene) and *UBC13* (Cre01.g046850) and present in the co-expression cohort linked to our modules. In addition, the gene *UBC22* (Cre12.g515450) appeared anti-correlated with other 26S proteasome subunit genes, hinting at a previously unexpected level of control.

We used the 96 genes that formed the protein degradation modules as baits to identify their co-expressed cohorts in each of our three most stringent networks (N1-N3). Via guilt-by association prediction, we thus assigned a potential function in protein degradation for 350-760 genes in addition to those already found within our modules (Supplemental Figure 10B, Supplemental Data Set 13).

#### Cilia Modules

Four modules were associated with GO terms with a function in cilia assembly or intraciliary transport. They also demonstrated partial overlap between themselves, indicating that these four modules defined a single, larger cilia group consisting of 221 nuclear genes (Figure 6F). The genes making up these modules were highly co-expressed with a fraction of genes identified in CiliaCut and the cilium proteome (Figure 6F). The intersection of the initial gene lists (modules, CiliaCut, overlap and cilium proteome) defined a set of 44 genes, nine of which (*ODA1*, *DRC3*, *IFT121*, *IFT46*, *IFT74*, *MBO2*, *MIA1*, *PF16* and *PF20*) were previously identified through forward genetic screens. We also extracted the co-expression cohorts associated with cilia modules, CiliaCut and the cilium proteome (Figure 6G-I, Supplemental data Set 8), linking several hundred genes to cilia. Their overlap (when using the N1 network) consisted of a set of 193 high-confidence cilia-related genes.

#### Photosynthesis modules

Four modules defined a larger photosynthesis group (Figure 6K) that we sub-divided into three modules containing many of the genes encoding tetrapyrrole biosynthetic enzymes, while the last module was related to photosystems components. We extracted their co-expression cohorts (Figure 6L-N), resulting in hundreds of genes exhibiting strong co-expression. We also determined the overlap between the initial gene lists (Figure 6O) and their N1 cohorts (Figure 6P): the co-expression modules clearly included both photosynthesis- and tetrapyrrole biosynthesis-related genes. As might be expected for genes necessary for proper chloroplast function, the overlap between N1 cohorts was substantial across all categories tested (modules, photosynthesis and tetrapyrroles), highlighting interesting genes for potential follow-up studies within the modules and the N1 cohort.

### Genes in Co-Expression Modules Cluster Based on their Diurnal Phase

During our analysis of co-expression modules, we noticed a high proportion of diurnal synchronization between co-expressed genes within modules and their associated co-expression cohorts, even though diurnally-expressed genes occupy the entire diurnal time landscape (Figure 7A, 7B). We therefore asked how frequently genes within co-expressed modules shared the same phase. Out of 117 modules extracted from the N3 network, 110 contained at least two rhythmic genes (Figure 7C). with a mean percentage of rhythmic genes of 65% and a median value of 71.6% (Figure 7C). Modules with few rhythmic genes tended to be associated with large standard deviations, indicative of little synchronization between the genes comprising them (Figure 7C). By contrast, modules consisting of a higher frequency of rhythmic genes showed high synchrony; their mean phase provided information relating to the biological function of each module, as illustrated below. Notably, the anti-correlated cohorts to most modules exhibited a mean phase that was 6-12 h out of phase with that of their related module (not shown), highlighting the importance of time-of-day when considering co-expression.

**Figure 7.**
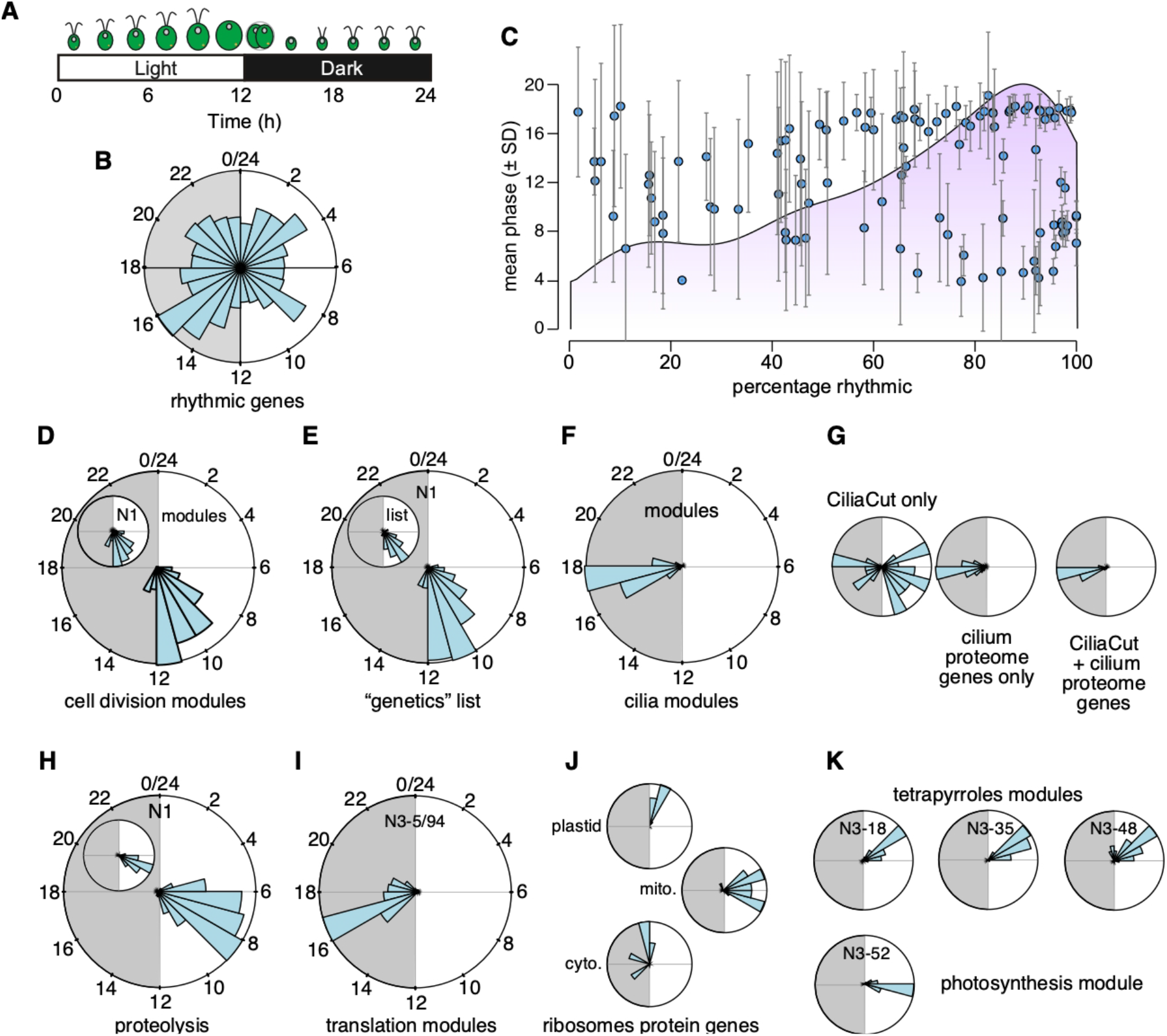
Co-Expression Modules Routinely Comprise Genes with Similar Diurnal Phases. **(A)** Schematic of the Chlamydomonas diurnal cycle in cell division events. **(B)** Phase distribution of 10,294 high-confidence diurnally rhythmic genes, shown as a circular plot covering the full 24 h of a complete diurnal cycle. Gray shade indicates night. **(C)** Co-expression modules with a high percentage of rhythmic genes exhibit a uniform diurnal phase. The light purple shade indicates the distribution of rhythmic modules. **(D-K)** Example of phase distribution for co-expression modules and associated N1 co-expression cohorts.

Molecular events leading to cell division are coordinately expressed with a phase distribution between 10-12 h after dawn: in agreement, we determined that the phase distribution of cell division modules and genes from the cell division “genetics” list showed the same phase preference, with 232 out of 245 genes being rhythmic, as did their associated co-expressed cohorts from the N1 network, (Figure 7D, 7E). After cell division, cells reassemble cilia in anticipation of the coming dawn: 191 (out of 221) genes within cilia modules exhibited a marked preference for the middle of the night part of the diurnal cycle, which precisely corresponds to the time of cilia biogenesis (Figure 7F). The degree of synchrony may provide an additional selection criterion for co-expressed genes, as seen with phase distributions of genes belonging to CiliaCut only (that is, CiliaCut genes whose gene products were not detected in the cilium proteome). Indeed, CiliaCut only genes displayed a wide range of diurnal phases, whereas co-expressed cilium proteome genes and genes at the intersection of CiliaCut and the cilium proteome were highly rhythmic and synchronized to the middle of the night (Figure 3C and Figure 7G).

We used the 96 genes (Figure 7H, inset) that form the protein degradation modules as baits to identify their co-expressed cohorts. They displayed a high degree of synchronized rhythmicity across diurnal datasets (Figure 7H). Only two out of the 96 genes from the protein degradation modules did not show rhythmic expression over a diurnal cycle. The occurrence of diurnal rhythmicity remained very high in their associated co-expression cohorts, with 391 rhythmic genes out of 450. The distribution of their diurnal phases was also quite narrow for both sets of genes, with a peak in the second half of the day (Figure 7H). We speculate that timed protein degradation offers a mechanism for the removal of photo-oxidized proteins, which is broadly consistent with the recent characterization of Chlamydomonas mutants lacking activities for the E3 ubiquitin ligase and Cullin components of the SCF (Skip, Cullin, F-box) complex (Gabilly et al., 2019).

The majority of genes that belonged to the non-redundant translation modules N3-5/94 were rhythmic (121 out of 158), and their diurnal phases concentrated in a narrow window of time between 3 and 5 h into the dark part of the diurnal cycle (Figure 7I). GO enrichment analysis indicated a role for these two modules in the nucleolus and ribosome biogenesis (Supplemental Table 3). Cytosolic *RPGs* were constitutively expressed and thus had no clear diurnal phase, whereas both plastid and mitochondrial *RPGs* exhibited preferred diurnal phases between 1-2 h and 3-5 h after dawn, respectively (Figure 7J), as expected (Zones et al., 2015).

Four modules defined a larger photosynthesis group that we sub-divided into three modules containing many of the genes encoding tetrapyrrole biosynthetic enzymes, while the last module was related to photosystems components. Both sub-groups were highly rhythmic over the diurnal cycle and restricted to a small time-window. Their respective phases agreed with their underlying biological function: genes encoding tetrapyrrole biosynthetic enzymes peaked ~ 2 h prior to components of both photosystems (Figure 6K). While highly co-expressed, photosynthesis- and tetrapyrroles-related modules did not substantially overlap (Supplemental Data Set 14), indicating that a diurnal phase difference of 2 h was sufficient to form independent clusters.

We conclude that co-expression modules are strongly influenced by the diurnal phase of their constituent genes. While this result may in itself not be surprising, it also raised the question of the overlap contribution of diurnal phase to clustering in our dataset, which we addressed next.

### Genes Cluster Based on their Diurnal Phase

While the majority of Chlamydomonas genes exhibit a diurnal expression profile when cells are grown under light-dark cycles, most of the samples included in our RNAseq dataset were collected from cells grown in constant light, with the assumption that cells in such cultures would be largely asynchronous. Since we observed frequent co-expression that followed diurnal phase information, we determined whether genes globally clustered according to their diurnal phase, and whether cells in constant light retained some entrained properties.

We first explored how various clustering methods ordered genes as a function of their diurnal phase. We performed this analysis on three datasets: the fully normalized and complete dataset (RNAseq4), which included samples collected from cells grown in constant light and under diurnal cycles; RNAseq4LL, only consisting of samples collected from cells grown in constant light; RNAseq4LD, comprising all samples with a rhythmic component, either diurnal or related to cell cycle progression. We calculated all pairwise PCCs and ordered genes according to hierarchical clustering (hclust, as shown in Supplemental Figure 5B), Angle of the Eigenvectors (AOE, Figure 8A), or First Principle Component (FPC, Supplemental Figure 11). The AOE correlation matrix exhibited a smooth transition from the first gene to the last gene (along each row), with strong positive correlations along the diagonal and at the upper right corner, separated by a gradual transition to negative correlations parallel to the diagonal (Figure 8A). The matrix also lacked the localized clustering seen with the “hclust” method (compare Figure 8A with Supplemental Figure 5B). The FPC correlation matrix arranged pairwise PCCs in a similarly smooth pattern, with the strongest positive PCC values located in the upper left corner and the strongest negative PCCs in the upper right corner (Supplemental Figure 11A). The PCCs generated from RNAseq4LD followed a wider normal distribution relative to those of RNAseq4 and RNAseq4LL (Figure 8B), which we hypothesize results from the smaller number of samples and a higher amplitude in gene expression under rhythmic conditions, in contrast to averaged values from asynchronous cells.

**Figure 8.**
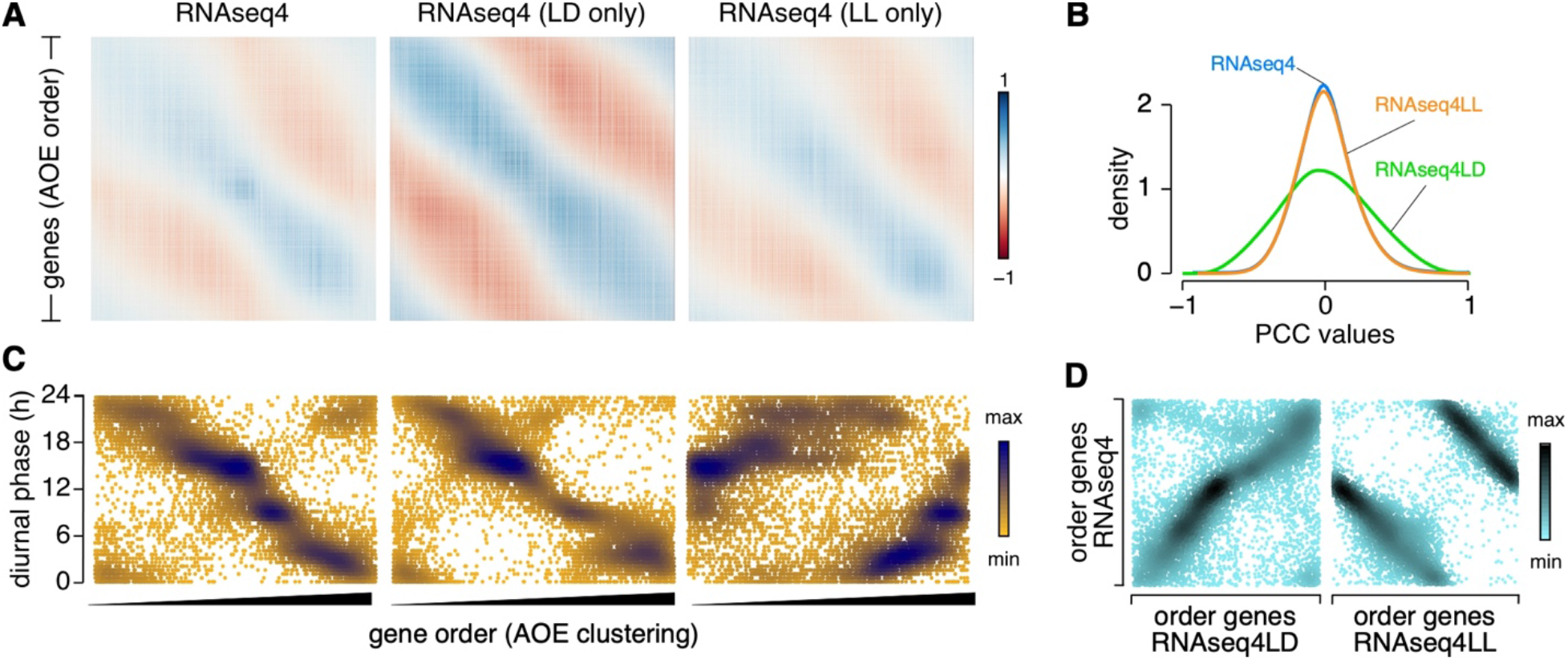
Genes Cluster Based on their Diurnal Phase. **(A)** Correlation matrix of the 17,741 Chlamydomonas nuclear genes, ordered based on clustering by the Angle of the Eigenvector (AOE) method built into *corrplot*, using the fully normalized dataset RNAseq4, RNAseq4LD (consisting of RNA samples collected from cells grown under light-dark cycles) and RNAseq4LL (with all other RNAseq samples) as input. **(B)** Distribution of pairwise PCCs for all gene pairs using RNAseq4, RNAseq4LD and RNAseq4LL as input. **(C)** Scatterplot of diurnal phases from 10,294 high-confidence diurnally rhythmic genes, as a function of their order from AOE clustering, using RNAseq4, RNAseq4LD and RNAseq4LL as input. We saved gene order following AOE clustering (from 1 to 17,741) and plotted the diurnal phase of the subset of 10,294 rhythmic genes (along the y axis). **(D)** Scatterplot of diurnal phases from 10,294 high-confidence diurnally rhythmic genes, ordered based on the AOE clustering method on RNAseq4 (y axis) and RNAseq4LD or RNAseq4LL (x-axis).

We next assigned a row number to each gene according to its place within the AOE correlation matrices, from 1 to 17,741. For those that also exhibited a diurnal expression pattern (Supplemental Figure 9; Zones et al., 2015, Strenkert et al., 2019), we plotted their diurnal phase (on the y axis) as a function of AOE gene order (on the x axis). As shown in Figure 8C, the relationship between AOE gene order and diurnal phases was far from random, and instead followed a linear pattern, whereby genes that appeared first in the AOE correlation matrix had phases with peaks in the late evening. As gene row number increased, diurnal phases gradually decreased, demonstrating the widespread influence of diurnal phase on correlation potential between gene pairs. In addition, the overall pattern of the AOE correlation matrix was reminiscent of that seen for diurnal experiments (Figure 1C, 1E), with genes separated by 12 h in terms of diurnal phases showing the strongest anti-correlations, while genes in similar time neighborhoods shared strong co-expression.

The RNAseq4 and RNAseq4LD datasets globally resulted in the same gene order after AOE clustering (Figure 8C), which at first might imply that samples collected from diurnally-grown cells imposed the observed gene ordering. However, this did not appear to be the case, as 1) the overall pattern of the AOE matrix for RNAseq4LL-derived PCC values was identical to that of RNAseq4 (Figure 8A), and 2) the corresponding gene order still carried diurnal information, as evidenced by the increase in diurnal phase with increasing gene order (Figure 8C), and despite the removal of all diurnal samples. Although the AOE clustering gene order did change between the RNAseq4 and RNAseq4LL matrices, the alteration in the pattern was systematic: a scatterplot of gene order for RNAseq4 and RNAseq4LL underscored the linear relationship between the two gene order series (Figure 8D). FPC clustering also sorted genes according to their diurnal phase, although along distinct parameters Supplemental Figure 11B).

We conclude that diurnal phase contributes substantially to the clustering of genes, even for samples obtained from cells grown in constant light. Such samples appear to retain diurnal information that shapes the clustering outcome at the genome level.

### Molecular Timetable Analysis Confirms Residual Synchronization of the Chlamydomonas Transcriptome

That genes clearly clustered according to their diurnal phases even in a dataset comprised solely of samples collected from cells grown in constant light raised the possibility that these samples exhibited residual rhythmicity. We thus applied the molecular timetable method (Ueda et al., 2004) to all RNAseq samples to determine the extent of rhythmicity they might exhibit. The molecular timetable method, whose principle is briefly explained in Supplemental Figure 12, extracts the rhythmic (diurnal or circadian) information from single time-point transcriptomes using the known phases and expected expression levels from a reference diurnal (or circadian) dataset. We selected 480 genes across 24 phase bins; their peak time of expression is known exactly, as well as their expression levels. We then extracted their normalized expression from RNAseq4 and calculated the mean expression for each phase bin. Finally, we plotted this mean for each RNAseq sample and each diurnal phase bin as a heatmap.

We first looked at the two large diurnal time-courses, shown in Figure 9A, to validate out methodology. Indeed, each diurnal sample (one row) showed a rhythmic pattern with each peak and trough separated by ~12 h. In addition, successive time-points were more similar to one another than to later time-points, as observed earlier in the correlation matrix (Figure 1E). These results demonstrated the applicability of the molecular timetable method to Chlamydomonas RNAseq samples, paving the way for the extraction of the internal time of the collected sample, as determined by the phase bin with maximal normalized expression.

**Figure 9.**
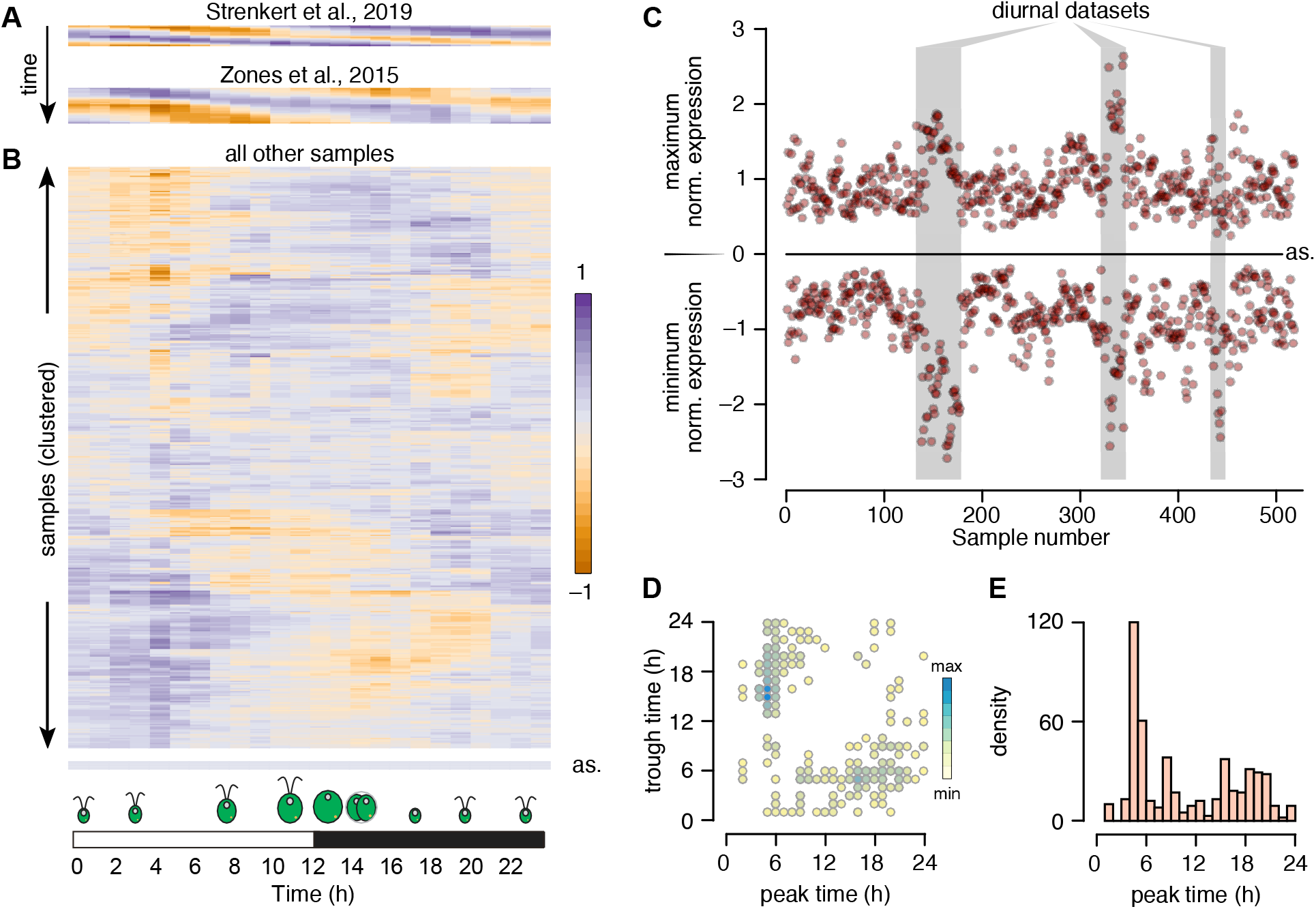
Chlamydomonas Cultures Grown in Constant Light Retain Substantial Rhythmicity. Heatmap representation of the molecular timetable approach, applied to two diurnal datasets: Strenkert et al., (2019) and Zones et al., (2015) **(A)**, and to all remaining RNAseq samples **(B)**. Each sample is represented as the mean expression of 20 phase marker genes (per h). In **(A)**, diurnal samples are ordered from top to bottom. For **(B)**, samples were subjected to hierarchical clustering while generating the heatmap in R. as: heatmap from an asynchronous sample, corresponding to the average expression of all rhythmic genes for each time-point. **(C)** Scatterplot of minimum and maximum normalized expression across all RNAseq samples. Diurnal time-courses are indicated by a gray shade. as: expected position of minima and maxima for a completely asynchronous sample. The samples are ordered by experiments: therefore, consecutive data points belong to the same experiment. **(D)** Peak and trough times largely occur 12 h apart. Scatterplot of all peak expression time (x-axis) and trough times (y-axis). **(E)** Distribution of peak times across all RNAseq samples.

We next subjected all remaining RNAseq samples to the same analysis and clustered them based on their underlying pattern while generating the heatmap shown in Figure 9B. Completely asynchronous samples should appear off-white across all phase bins (“as”, bottom of Figure 9B); overwhelmingly, Chlamydomonas RNAseq samples instead displayed remarkable residual rhythmicity. Diurnal time-courses were easy to distinguish from other samples when we plotted the minimum and maximum normalized expression values associated with each sample (Figure 9C). Notably, most other samples, collected from cells grown in constant light, retained strong global oscillations, which we estimate to represent a synchronization between cells ranging from 21-96%, with a mean rhythmicity of 48%, based on the amplitude between minima and maxima relative to diurnal time-course samples (Figure 9C).

The timing of minimum and maximum gene expression should be ~ 12 h apart in diurnal and rhythmic samples: we therefore plotted peak and trough times predicted for all samples based on the molecular timetable data. As shown in Figure 9D, most samples indeed reached peak value 12 h after their lowest time-point, validating our hypothesis that the vast majority of Chlamydomonas RNAseq samples exhibit strong residual rhythmicity even when the cells were grown in constant light.

Finally, we asked whether samples displayed a preferential diurnal phase by plotting the distribution of peak phases across all samples. To our surprise, about one third of all samples showed a peak phase between 5-6 h after dawn.

### Applicability of the Molecular Timetable Method to Other Algae: *Volvox carteri* and *Chromochloris zofingiensis* as Tests

Incorporating new Chlamydomonas transcriptome datasets to the one we used here would be cumbersome, as it would entail repeating all normalization steps each time a new dataset is added. A more practical approach would be to subject new transcriptome datasets to an abridged normalization, namely log_2_ normalization followed by normalization to the mean calculated from our full dataset. We tested the usefulness of this method by re-analyzing a transcriptome dataset included in our original list that was focused on iron homeostasis (Urzica et al., 2012c), for which Chlamydomonas cells had been grown with various iron concentrations (0.25, 1 or 20 μM FeEDTA) in autotrophic (no reduced carbon source provided, but cultures were bubbled with CO_2_) or heterotrophic (with acetate as reduced carbon source) conditions. We normalized FPKM counts to the mean inferred from the full RNA-seq dataset, and used the same diurnal phase values as above. As shown in Figure 10A, autotrophic cultures exhibited a very similar molecular timetable profile, with an estimated internal phase around dawn across all three iron concentrations. In sharp contrast, heterotrophic cultures responded very differently: indeed, iron-limited cultures (0.25 μM FeEDTA) were 12 h out of phase with the other two samples. Iron-limited heterotrophic cultures grow more slowly than iron-deficient (1 μM FeEDTA) or iron-replete cultures (20 μM FeEDTA). We hypothesize that the difference in internal phase between heterotrophic samples may thus partially reflect the time at which cultures were sampled, as cells were harvested at the same cell density (Urzica et al., 2012c). However, we cannot exclude a contribution to a slower circadian clock under low iron conditions, as described for land plants (Chen et al., 2013; Salomé et al., 2013; Hong et al., 2013). Nonetheless, we conclude that the molecular timetable method is applicable to Chlamydomonas samples after performing log_2_ and mean normalization.

**Figure 10.**
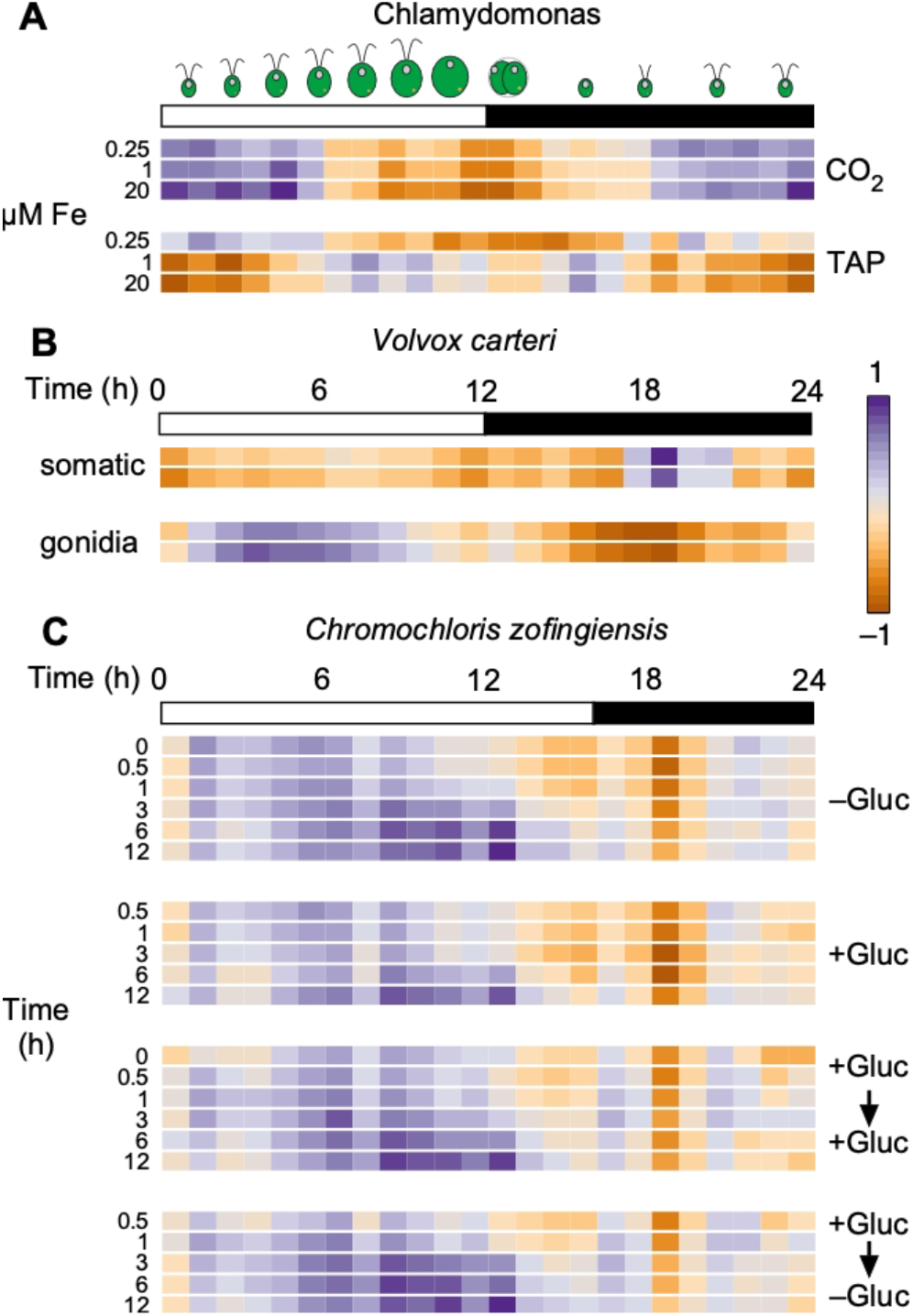
Application of the Molecular Timetable Method to Independent RNAseq Experiments Across Algae. **(A)** Re-analysis of a transcriptome dataset including in our initial RNAseq data (Urzica et a., 2012). We subbjected FPKM values to log_2_ normalization, followed by normalization to the mean (obtained during the normalization steps that yielded RNAseq4). We then used the molecular timetable method to determine the rhythmic pattern of the samples (Chlamydomonas CC-4532 strain grown in Tris Acetate Phosphate (TAP) or Tris Phosphate (CO_2_) medium with 0.25, 1 or 20 μM FeEDTA. **(B)** Molecular timetable method applied to *Vovox carteri* samples collected in duplicates from somatic or gonidial cells (Matt and Umen, 2018). **(C)** Molecular timetable method applied to *Chromochloris zofingiensis* samples collected over 12 h after addition and remval of glucose (Roth, Gallaher et al., 2019). For **(A)**, we used 960 highly rhythmic genes to draw the heatmap. For **(B)** and **(C)**, we included all rhythmic genes with orthologs in *V. cateri* **(B)** or *C. zofingiensis* **(C)**, after log_2_ normalization and normalization with the Chlamydomonas-derived gene means.

We then explored the applicability of this method to other algae where a high-density diurnal time course is not available: *Vovox carteri* and *Chromochloris zofingiensis*. *V. carteri* samples consisted of two technical replicates each collected from somatic and gonidial cells (Matt and Umen, 2018). We obtained one-to-one orthologs between Chlamydomonas and *V. carteri* from Phytozome, after which we subjected all *C. carteri* genes with a rhythmic Chlamydomonas ortholog to log_2_ normalization and to normalization with Chlamydomonas means. We then calculated the average normalized expression for all genes, in 1 h bins. Gonidial cells appeared strongly rhythmic, with a peak phase around 4-5 h after dawn and a trough ~12 h later (Figure 10B). Remarkably, somatic cells exhibited a completely different profile with a peak phase in the middle of the night. We performed the same analysis of transcriptome samples collected in *C. zofingiensis* over a 12 h time-course with addition or removal of glucose from the growth medium (Roth et al., 2019). Here, cultures were maintained in light-dark cycles consisting 16 h light and 8 h darkness. All samples exhibited a rhythmic profile, strongly indicating that the molecular timetable accurately predicted the internal phase of the samples. Indeed, the peak phase of samples collected later during the day showed a clear and distinct shift to a later phase. Notably, the rhythmic pattern extracted from these transcriptome samples followed the same overall pattern regardless of the treatment imposed on the cultures, which is consistent with the strong contribution of time-of-day noted in these samples (Roth et al., 2019).

We conclude that the molecular timetable method can be applied to Chlamydomonas and to other algae, even when they lack a reference diurnal time-course. Such analysis would allow a rapid estimation of the contribution of rhythmic gene expression to variation in gene expression, even in the absence of a reference diurnal time-course.

## DISCUSSION

We initially set out to analyze multiple RNAseq datasets to prioritize genes whose expression responded to changes in iron status in Chlamydomonas and Arabidopsis. Our working hypothesis was that genes with a prominent role in iron homeostasis should closely follow the expression pattern of known iron-responsive genes like the iron transporters *IRT1* and *IRT2* or *NATURAL RESISTANCE ASSOCIATED MACROPHAGE PROTEIN 4* (*NRAMP4*), the *Fe ASSIMILATION* (*FEA*) genes *FEA1* and *FEA2*, or the *FERRIC REDUCTASE* (*FRE*). We quickly realized that the assembly of 518 RNAseq samples into one dataset offered a unique opportunity to explore the transcriptome landscape of the alga. We believe that we have only skimmed the surface during our meta-analysis and invite others to use this dataset for their own research questions.

We were surprised to see how little correlation existed between Chlamydomonas experiments, even though several queried the same biological question, such as responses to nitrogen deficiency or metal deficiencies (Figure 2). Samples collected in the same laboratory similarly failed to show strong correlations, although growth conditions are likely to be similar. We do not fully understand the underlying source of variation, but we propose that strong residual rhythmic gene expression may contribute to the observed pattern. As a test of our analysis pipeline, we determined the correlation matrix of Arabidopsis microarray datasets, downloaded from AtGenExpress. As shown in Supplemental Figure 13, samples (using the expression data for all genes as data points) clearly grouped as a function of the tissue of origin, with shoot and leaf samples generally strongly correlated, while anti-correlated with root samples. It is likely that Arabidopsis samples show strong differentiation of their expression profiles as a function of the tissue of origin, as might be expected, thus validating our pipeline. More puzzling though is the fact that Chlamydomonas samples behave as independent units that share little correlation with others. In this regard, it would be informative to perform a comparative analysis of transcriptome datasets from multiple uni- and multi-cellular organisms, to determine whether multicellularity drives the more polarized differentiation of expression profiles seen across Arabidopsis samples relative to Chlamydomonas.

Co-expression modules assemble the most consistent gene pairs into a coherent list that is characterized by high connectivity. However, each gene is itself co-expressed with many genes not included in the module (Supplemental Figure 8). These co-expression cohorts can provide cues as to the function of a gene, especially when it does not belong to a module. In addition, genes with the opposite expression profile can give hints as to the function of a gene of interest. We have extracted co-expression and anti-correlation cohorts for all Chlamydomonas genes, provided as Supplemental Data Sets 2-7. We also provide an example script to run the same analyses presented here on any gene list, from extracting the co-expression cohort to plotting the corresponding correlation matrix (Supplemental File 1). We hope that this type of analysis spurs new discoveries, not only in Chlamydomonas but also in Arabidopsis and other plants. Our results with Arabidopsis *RPGs* (Figure 4E) demonstrates the applicability of the method to other organisms.

We do not anticipate all candidate genes identified based on co-expression to be functionally tested, at least not in Chlamydomonas. Rather, we expect co-expressed genes to be compared to other gene lists, generated by other means, in order to narrow down the number of interesting candidates for follow-up studies further. For example, large-scale non-targeted mutant screens in Chlamydomonas pave the way for the systematic genetic dissection of phenotypes (Li et al., 2015, 2019); we envisage that the intersection of co-expression and large-scale genetic screens will empower research, not only in Chlamydomonas, but also in other algae.

The Chlamydomonas life cycle resolves around cell division, the timing of which can be synchronized to dusk by light-dark cycles (Zones et al., 2015; Strenkert et al., 2019; Cross and Umen, 2015). When maintained under entraining conditions, at least 80% of the Chlamydomonas transcriptome exhibits rhythmic expression. It is unclear how quickly algal cells become asynchronous when transferred to constant light conditions. It is thought that cultures grown in constant light are largely arrhythmic at the population level due to loss of synchrony. When applying the molecular timetable to Chlamydomonas RNAseq samples, we discovered that the vast majority of samples exhibited substantial rhythmicity, even when collected from cells grown in constant light (Figure 9). About one third of all samples appeared to have been collected 5-6 h after subjective dawn (that is, the dark-to light transition, had the cells been maintained under entraining conditions). Based on the amplitude between minima and maxima extracted from phase marker genes, we estimate that 21-96% of cells within a given culture were synchronized, with a mean of 48%. Chlamydomonas strain stocks are typically kept in constant light on solid medium before inoculating a liquid culture, which will itself be placed in constant light. Pre-cultures are common before inoculating the test culture; cells are generally collected by centrifugation when they reach mid-log. It is therefore possible that diluting cells at the beginning of an experiment sends a resetting signal to the Chlamydomonas circadian clock, the signature of which is still present 2-3 d later, as evidenced by the degree of residual synchronization in all samples analyzed. We are here only seeing the bulk behavior of Chlamydomonas cultures. Single-cell RNAseq (scRNA-seq) analysis will allow a more detailed dissection of the diurnal contribution to the Chlamydomonas transcriptome landscape. To begin to explore this possibility, we recently performed scRNA-seq on almost 60,000 Chlamydomonas cells grown under three growth conditions and from two genotypes. Indeed, we observed a substantial heterogeneity among the cells that was partially explained by the endogenous phase of individual cells (Ma et al.). Although cultures were grown in constant light for several weeks, we hypothesize that diluting cells at the beginning of an experiment may act as a resetting signal for the endogenous cell cycle and circadian clock.

Our observations also raise a question regarding the design of RNA-seq experiments: when assessing the effect of a mutation or a treatment on cultures, Is it more important to collect samples at the same cell density or at the same time? Our results suggest that sampling time exerts a far greater influence on expression outcomes than sampling density would. Best practices for RNAseq analysis may therefore dictate that a matched control sample be collected at each time-point in order to remove any contribution to differential gene expression from the strong rhythmicity exhibited by cultures. Genes belonging to the same co-expressed modules tended to have the same diurnal phase (Figure 9C); the narrow window of expression seen in rhythmic genes would thus be missed when comparing samples collected hours apart. In Arabidopsis, samples collected 30 min apart already exhibited differential expression (Hsu and Harmer, 2012). Our results generalize this observation.

The molecular timetable method is a powerful and easily implemented method to test the rhythmic component of transcriptome data. We demonstrate here that Chlamydomonas data can be transferred onto other algae like *V. carteri* and *C. zofingiensis* to reveal an unexpected dimension of rhythmic expression from single time points. We propose that all transcriptome datasets should be subjected to such analysis before delving into more in depth analysis, to estimate the fraction of variation in gene expression that might be explained by rhythmic expression. We provide the mean and phase values from Chlamydomonas to normalize RNAseq data from other algae as Supplemental Data Set 15.

In conclusion, we describe here an analysis of co-expression in the green unicellular alga Chlamydomonas. We observed known and new connections between genes ad provide the tools to take this analysis further for any gene of interest, in both Chlamydomonas and other system with a body of transcriptome data available.

## MATERIALS AND METHODS

### Co-Expression Analysis Network in Chlamydomonas

We re-analyzed a set of 58 RNAseq experiments, consisting of 518 samples, by mapping reads to version v5.5 of the Chlamydomonas genome (v5.5 from Phytozome) with STAR (v2.5) (Dobin et al., 2013) using default settings except --alignIntronMax 10000 --outFilterMismatchNoverLmax 0.04. Expression was calculated in terms of Fragments Per KB per Million mapped reads (FPKMs) with cuffdiff (v2.0.2) (Trapnell et al., 2014) using default settings except --multi-read-correct --max-bundle-frags 1000000000. We assembled all expression estimates as FPKM into one file and did not attempt to correct for batch effect at this stage, with the thought that such effects would contribute to the variation in expression. We then log_2_-transformed mean FPKMs across replicates was with a pseudo-count of “1” added prior to conversion, followed by quantile normalization with the R package *preprocessCore*. Finally, we subtracted mean expression across all experiments for each gene, which removed any potential batch effects from the data. We calculated Pearson’s correlation coefficients (PCC) with the *cor* function in R and visualized for each gene pair using the R package *corrplot*, using all 518 expression estimates. We maintained four expression datasets following each normalization step: RNAseq1 (mean FPKMs); RNAseq2 (log_2_-normalized); RNAseq3 (quantile-normalized); RNAseq4 (normalized to mean).

We calculated the rank for all gene pairs by inverting the sign of PCCs by multiplying the data frame by –1, then converting PCC values for each gene into ranks with the function *rank* in R. We derived the mutual ranks (MRs) for two genes a and b from the formula MR(a,b) = √(rank _a→b_ × rank _b→a_). Considering a matrix of ranks, the ranks rank_a⟶b_ and rank_b⟶a_ are geometrically linked on either side of the diagonal: if rank_a⟶b_ has the coordinates (x,y) in the rank matrix, then rank_b⟶a_ will have the coordinates (y,x). We therefore transposed the rank matrix with the *t* function in R. We obtained MR values for each gene pair by multiplying each cell from the rank matrix by their counterpart in the transposed rank matrix, then square-rooted.

For network selection and visualization, we calculated edge weights from MR values with the formula: Nx = e^−(MR-1)/x^, with x = 5, 10, 25, 50 or 100. Only Nx ≥ 0.01 were considered significant. We extracted gene pairs with significant edge weights from the full edge weight matrix and loaded them into Cytoscape 3.5.1. We detected modules of co-expressed genes with ClusterONE with default parameters. Modules with a p-value ≤ 0.1 were considered significant.

We also determined lists of anti-correlated genes by ranking PCC values from the non-inverted PCC matrix generated by *corrplot*, and by calculating associated edge weights as above. In this case, we limited our analysis to identifying anti-correlated genes, as ClusterONE cannot detect modules using edge weights from anti-correlated genes.

### Co-expression Analysis Network in Arabidopsis

Microarray datasets were downloaded from the AtGenExpress project site (http://jsp.weigelworld.org/AtGenExpress/resources/), and collated into a single file that consisted of 34 Arabidopsis accessions, 16 sets of etiolated seedlings exposed to various light treatments, 36 sets of seedlings exposed to pathogens, 13 cell culture samples, 68 sets each for shoots and roots exposed to various abiotic stresses, 79 developmental samples (72 from shoots or leaves, 7 from roots), and 18 sets each for leaves and roots subjected to iron deficiency, with controls included. We log_2_-normalized all data when not already done, and followed the same normalization steps described for the Chlamydomonas data set.

### Analysis of Co-Expression from ClusterONE Modules

We extracted normalized expression data (from RNAseq4) for genes belonging to a given cluster in R using the *stack* and *unstack* functions, and generated the corresponding co-expression matrix with *corrplot*. We tested for overlap between co-expression modules with similar predicted function with the online tool Venny (Oliveros, 2007), and redrew co-expression matrices with a non-redundant gene list as input. Unless stated otherwise, we ordered genes based on the FPC (First Principle Component) clustering method built into *corrplot*.

### Analysis of Co-expression from Manually-Curated and Community Gene Lists

We extracted normalized expression data for genes that belonged to manually-curated or community-generated lists as described above for co-expression modules. We maintained the same gene order when working with community lists, as the genes were sorted and grouped based on shared function. We sorted genes from manually-curated lists following the FPC method in *corrplot*.

### Identification of Co-Expression Cohorts

We extracted the sets of genes co-expressed with each gene belonging to our co-expression modules in R by merging each module-specific gene list with a file representing all nodes and edges from networks N1 to N3. We collapsed each co-expression cohort into a non-redundant list by using the function *unique* in R and tested each subset for overlap with *merge* or *join*.

Manually-curated and community-generated gene lists presented an initial challenge, since not all of their constituents are necessarily co-expressed (for example, only a fraction of the genes defined by the mutant screen carried out by Fred Cross for cell cycle mutants is co-expressed). We therefore 1) ordered genes using the FPC clustering method; 2) counted how many gene pair PCCs were above 0.25, 0.4 or 0.5 for each row of the matrix in order to 3) define cut-offs between subsets of genes with high-, medium- or low-PCCs. We then used these subsets (from 1 to 3) as bait to identify their associated co-expression cohort, as described above for co-expression modules.

### GO Category Enrichment in Co-Expressed Modules

We tested our co-expression modules for Gene Ontology term enrichment by using the PANTHER database (pantherdb.org) through the Gene Ontology Resource page (http://geneontology.org). First, all Chlamydomonas gene identifiers (Crexx.gxxxxxxx) were converted to their corresponding Uniprot identifiers using a gene-to-Uniprot list generated in-house. Of 117 modules, 86 retained at least 10 genes with corresponding Uniprot identifiers (31 had ≤ 9 genes with matching Uniprot identifiers and were deemed too small for further analysis), and 37 returned significant enrichment in GO term(s) for Biological Process.

### Venn Diagrams and Gene List Overlaps

We compared gene lists and determined the extent of overlap with the online tool Venny (Oliveros, 2007). Proportional Venn diagrams were drawn with BioVenn (Hulsen et al., 2008) for 2-way diagrams or EulerAPE 3.0.0 (Micallef and Rodgers, 2014) for 3-way diagrams.

### Statistics

PCC values for the entire genome were calculated with the *cor* function in R, and their distributions plotted with the *density* function in R. A random normal distribution of mean = 0 and standard deviation = 0.2 was generated with the *rnorm* function in R for 100 million values; only 23 values fell outside of the –1 to +1 range and were not discarded.

For comparisons between distributions, we applied a Kolmogorov-Smirnov test (ks-test) using the *ks.test* function in R.

## SUPPLEMENTAL MATERIALS

**Supplemental Figure 1**. Normalizations of the Chlamydomonas transcriptome dataset.

**Supplemental Figure 2**. How ribosomal protein genes (*RPGs*) respond to each normalization step.

**Supplemental Figure 3**. The R package *corrplot* and visualization of large correlation matrices.

**Supplemental Figure 4**. Correlations between experimental samples and normalization methods.

**Supplemental Figure 5**. Chlamydomonas gene pairs are largely not co-expressed.

**Supplemental Figure 6**. Co-expression cohorts for Chlamydomonas ferredoxins.

**Supplemental Figure 7**. From co-expression cohorts to co-expression modules.

**Supplemental Figure 8**. Using module nodes as baits to identify co-expressed genes.

**Supplemental Figure 9**. Convergence of diurnal phase between two time-courses.

**Supplemental Figure 10**. Co-expression of the protein degradation machinery is limited to the 26S proteasome.

**Supplemental Figure 11**. Genes Cluster Based on their Diurnal Phase.

**Supplemental Figure 12**. Molecular timetable method to extract diurnal information from single time-points.

**Supplemental Figure 13**. Arabidopsis microarray data clearly differentiates between tissue types.

**Supplemental Table 1**. Summary of expression estimates across all conditions and samples.

**Supplemental Table 2**. Cohort and modules sizes for co-expression data derived from the RNAseq4 dataset.

**Supplemental Table 3**. Summary of GO terms enriched in N3 co-expressed clusters.

**All Supplemental Data Sets have been uploaded to:** https://drive.google.com/drive/folders/1Ee9tArvYiMHgzx9fJ7-L-06xSc3uhPwj?usp=sharing

**Supplemental Data Set 1**. The fully normalized RNAseq dataset.

**Supplemental Data Set 2**. List of co-expressed genes for each nuclear Chlamydomonas gene for the N1 network.

**Supplemental Data Set 3**. List of co-expressed genes for each nuclear Chlamydomonas gene for the N2 network.

**Supplemental Data Set 4**. List of co-expressed genes for each nuclear Chlamydomonas gene for the N3 network.

**Supplemental Data Set 5**. List of anti-correlated genes for each nuclear Chlamydomonas gene for the N1 network.

**Supplemental Data Set 6**. List of anti-correlated genes for each nuclear Chlamydomonas gene for the N2 network.

**Supplemental Data Set 7**. List of anti-correlated genes for each nuclear Chlamydomonas gene for the N3 network.

**Supplemental Data Set 8**. Cilia genes, and co-expressed cohorts.

**Supplemental Data Set 9**. The fully normalized Arabidopsis dataset.

**Supplemental Data Set 10**. Co-expression cohorts for all *FDX* genes.

**Supplemental Data Set 11**. List of genes from the 117 co-expression modules identified in network N3.

**Supplemental Data Set 12**. Cell division modules and co-expressed cohorts.

**Supplemental Data Set 13**. Photosynthesis and tetrapyrroles biosynthetic genes and their co-expressed cohorts.

**Supplemental Data Set 14**. Proteasome and protein degradation-related genes and their co-expressed cohorts.

**Supplemental Data Set 15**. Phases and means for 10,294 rhythmic Chlamydomonas genes.

**Supplemental Data Set 16**. List of co-expressed genes for each nuclear Arabidopsis gene for the N1 network.

**Supplemental Data Set 17**. List of co-expressed genes for each nuclear Arabidopsis gene for the N2 network.

**Supplemental Data Set 18**. List of co-expressed genes for each nuclear Arabidopsis gene for the N3 network.

**Supplemental File 1**. Examplar R script to extract data for a gene list, plot the corresponding correlation matrix and extract the co-expression cohort.

**Supplemental File 2**. Normalization pipeline to turn transcriptome data into an input file for co-expression analysis.

## ACKNOWLEDGMENTS

Work in the Merchant laboratory is supported by a cooperative agreement with the US Department of Energy Office of Science, Office of Biological and Environmental Research program under Award DE-FC02-02ER63421. We thank Sean D. Gallaher and Ian K. Blaby for critical reading of the manuscript. We also acknowledge Michael Leonard for his efforts in remapping the Chlamydomonas transcriptome datasets used here.

## AUTHOR CONTRIBUTIONS

PAS designed and conducted all analyses with supervision from SSM. PAS wrote the manuscript with input from SSM.

